# Structure-Activity Relationships in the Design of Mitochondria-Targeted Peptide Therapeutics

**DOI:** 10.1101/2021.11.08.467832

**Authors:** Wayne Mitchell, Jeffrey D. Tamucci, Emery L. Ng, Shaoyi Liu, Hazel H. Szeto, Eric R. May, Andrei T. Alexandrescu, Nathan N. Alder

**Author notes:** Corresponding Author: Dr. Nathan N. Alder, 91 N. Eagleville Rd, Storrs, CT 06269-3125.

## Abstract

Mitochondria play a central role in metabolic homeostasis; hence, dysfunction of this organelle underpins the etiology of many heritable and aging-related diseases. Mitochondria-targeted tetrapeptides with alternating cationic and aromatic residues, such as SS-31 (Elamipretide), show promise as therapeutic compounds. In this study, we conducted a quantitative structure-activity analysis of three alternative tetrapeptide analogs that differed with respect to aromatic side chain composition and sequence register, benchmarked against SS-31. Using NMR and molecular dynamics approaches, we obtained the first structural models for this class of compounds, showing that all analogs except for SS-31 form compact reverse turn conformations in the membrane-bound state. All peptide analogs bound cardiolipin-containing membranes, yet they had significant differences in equilibrium binding behavior and membrane interactions. Notably, the analogs had markedly different effects on membrane surface charge, supporting a mechanism in which modulation of membrane electrostatics is a key feature of their mechanism of action. All peptide analogs preserved survival and energy metabolism more effectively than SS-31 in cell stress models. Within our peptide set, the analog containing tryptophan side chains, SPN10, had the strongest impact on most membrane properties and showed greatest efficacy in cell culture studies. Taken together, these results show that side chain composition and register strongly influence the activity of these mitochondria-targeted peptides. Furthermore, this work helps provide a framework for the rational design of next-generation therapeutics with enhanced potency.

## Introduction

As regulators of energy metabolism, mitochondria house the oxidative phosphorylation (OXPHOS) complexes that produce >90% of cellular ATP. Mitochondria also coordinate key cellular processes including lipid biosynthesis, ion homeostasis, and cell death. Consequently, mitochondrial dysfunction, particularly in tissues with high energy demand, is central to the etiology of many complex pathologies including cancer, cardiopathy, neurodegeneration, aging-related ailments, and heritable (primary) mitochondrial disease. Despite this, there are currently no FDA-approved therapeutics for the treatment of mitochondrial diseases.

Mitochondria-targeted cationic-aromatic tetrapeptides are among the most promising pharmacological interventions under development for the treatment of mitochondrial dysfunction. Also termed Szeto-Schiller (SS) peptides, these first-in-class compounds are synthetic C-terminally amidated tetrapeptides with a motif of alternating cationic and aromatic residues that is thought to be important for their ability to traverse membranes in a variety of cell types and concentrate in mitochondria (1–3). Many *in vitro*, preclinical, and clinical studies, predominantly with the lead compound SS-31 (Elamipretide), support the therapeutic efficacy of these peptides. Studies with isolated mitochondria and cell cultures show that SS-31 improves electron transfer efficiency and increases ATP production while reducing electron leak and reactive oxygen species (ROS) production (3–7). Furthermore, animal studies have demonstrated the ability of SS-31 to maintain cellular bioenergetics under stress conditions such as ischemia, hypoxia, and aging-related dysfunction (8–11). Finally, the clinical efficacy of SS-31 has been demonstrated for primary mitochondrial disorders (12, 13) and for age-related chronic diseases associated with mitochondrial dysfunction (14).

Progress toward elucidating the molecular mechanism of action (MoA) of these peptides has come on several fronts. Early studies suggested that SS peptides target the lipid bilayers of mitochondrial membranes through interactions with cardiolipin (CL) (4, 5, 15), the anionic phospholipid that is enriched in the inner mitochondrial membrane (IMM) and required for proper membrane morphology as well as function of membrane-bound complexes (16). This bilayer-mediated mechanism is supported by work with model systems in which peptide inhibited peroxidase activity of cytochrome *c* (4) and improved cristae ultrastructure (17). Recent work from our group quantitatively evaluated SS-31 interactions with CL-containing membranes, showing that the peptide affected lamellar bilayer properties (e.g. lipid lateral diffusion and packing interactions), with the most notable effect being on membrane electrostatics based on down-regulation of the surface potential (ψ_s_) that originates from the negatively-charged membrane interface (18). Other recent work has focused on the interactions of SS-31 with mitochondrial proteins. Based on a crosslinking/mass-spectrometry approach with a biotinylated SS-31 variant, the SS-31 interactome was shown to include a subset of membrane complexes primarily involved in ATP-generating processes (19). Moreover, in aged cardiomyocytes, SS-31 was shown to reduce proton leak mediated by the adenosine nucleotide transporter (ANT1) and stabilize the ATP synthasome (11). Notably, the vast majority of these SS-31-interactive proteins are known to bind CL, supporting a role of mitochondrial lipid composition in the molecular interactions of these compounds. Yet despite these insights, the lack of information relating the structure and function of these mitochondria-targeted tetrapeptides presents a barrier to a full understanding of their MoA.

An effective strategy to address this mechanistic knowledge gap is to test the effects of expanding the sequence space of mitochondria-targeted peptides using structure-activity analyses. In this study, we compared three sequence-variant peptide analogs that differed with respect to aromatic side chain content and cationic/aromatic register, using SS-31 as a benchmark. Our results provide a direct comparison of these tetrapeptide variants with respect to their membrane-bound conformations, effects on membrane properties, and relative efficacies in preserving cellular viability under stress. This work reveals that side chain composition has a profound effect on the structure-activity relationships of these mitochondria-targeted peptides. Most notably, the analog containing tryptophan side chains had the greatest potency in cell stress models, which we can correlate with its molecular-level interactions and effects on reductionist membrane systems. This work sets the foundation for the rational design of next-generation tetrapeptide variants with enhanced efficacy as mitochondrial therapeutics.

## Results

### Tetrapeptide analog set: design and rationale

In this study, we compared a test set of four tetrapeptides with different sequences (**Fig. 1A**). An alphabet of two basic residues (Arg, Lys) and three aromatic residues (Phe, Tyr, Trp) gives 3^2^ × 2^2^ × 2 = 72 possible sequence permutations with an alternating aromatic (φ) / basic (B) sequence periodicity (B-φ-B-φ or φ-B-φ-B). However, the number of sequences becomes much larger if D-amino acids (which can extend the medicinal lifetimes of peptides) and/or unnatural amino acids (which increase functional versatility) are included. A large library of peptides precludes detailed structural and functional studies, so we focused on a limited test set of peptides to investigate two fundamental properties: (i) the side chain register (B-φ-B-φ vs. φ-B-φ-B) and (ii) the types of aromatic side chains. The cationic/aromatic register has potential structural ramifications, for example, in determining the polar interactions between peptide basic groups and CL. Aromatic amino acid type can modulate hydrophobicity, aromaticity, polarity, and hydrogen bonding capacities, which in turn can affect both peptide structure and peptide-membrane interactions (20).

**Figure 1.**
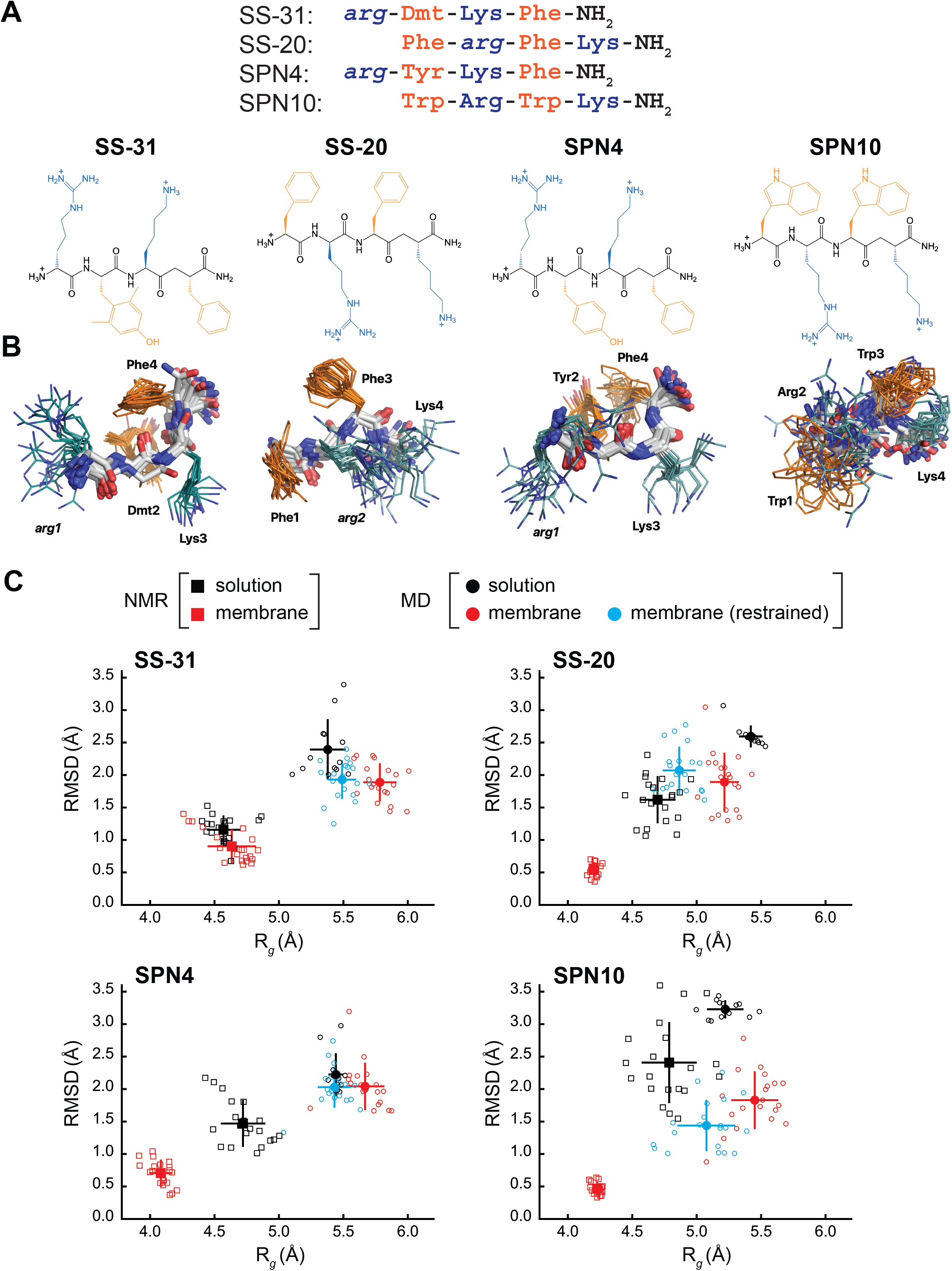
Peptide structures and membrane-driven peptide folding. **A) Peptide sequences and chemical structures.** *Upper series*, peptide sequences aligned with respect to basic and aromatic side chains; *lower series*, chemical structures of peptides. Basic and aromatic groups are shown in teal and orange, respectively. **B) NMR structures of peptides in solution.** The top 20 lowest-energy NMR conformers obtained by ROESY analysis. The main chain atoms are colored by type (carbon, white; oxygen, red; nitrogen, blue); basic and aromatic side chains are colored in teal and orange, respectively. **C) Comparison of peptide conformational variance in NMR and MD assemblies.** Radius of gyration (R_*g*_) and root mean square deviation (RMSD) to the average peptide structure calculated for NMR structures in solution (*black squares*) and in the presence of a membrane (*red squares*), and for MD simulations of peptides in solution (*black circles*), in the presence of a lipid bilayer conducted without (*red circles*), and with (*cyan circles*) NOE restraints. Measurements of individual peptides (n=20) are shown as small symbols, with ensemble averages and error bars (SD) shown as larger symbols. Corresponding values for the R_*g*_ and RMSD measurements are in Supplementary Table S4.

Our test set (**Fig. 1A**) included two analogs with B-φ-B-φ register, SS-31 (our benchmark) and SPN4. These two analogs differ only with respect to the second-position aromatic residue. SS-31 contains the unnatural amino acid 2,6-dimethyltyrosine (Dmt), known to be important for free radical scavenging by this peptide (1, 3, 21). By contrast, SPN4 replaces this Dmt with Tyr. Proteinogenic Tyr retains the phenolic OH group that can scavenge radicals and mediate H-bond interactions but allows us to evaluate the effect of the two tyrosine methyl groups on peptide structure and function. Our test set also included two analogs with φ-B-φ-B register, SS-20 and SPN10. With its Phe/Phe aromatic composition, SS-20 does not possess the free radical scavenging properties of SS-31; however, it has also demonstrated efficacy with many mitochondrial disease models (22–24), confirming that scavenging activity is not an essential feature of the MoA of this class of compounds. Finally, SPN10 is unique among our test set in that it contains only L-enantiomer side chains and it has two Trp residues, which contain the bulky bicyclic indole group with a pyrrole-like NH that can mediate H-bond interactions.

### The free peptides are extended but have residual structure due to aromatic interactions

As a first step toward comparing the four peptides, we determined their NMR structures in solution. Small water-soluble and membrane-active peptides typically adopt their bioactive conformations only upon binding membranes (25, 26). However, even very short peptides can have preferred conformations in aqueous solution (27), particularly if they are enriched in aromatic residues. Structural analysis of these peptides in solution can shed light on their properties in the extracellular milieu, the cytosol, and aqueous mitochondrial subcompartments.

NMR spectra of the peptides were assigned using 2D TOCSY and NOESY spectra for ^1^H signals, and ^1^H-^13^C HSQC and ^1^H-^15^N HSQC experiments with samples at natural isotope abundance for ^13^C and ^15^N nuclei, respectively (**Table S1**). The free peptides in solution show few, if any, NOEs (**Fig. S1A**), which is consistent with their molecular masses of ∼600 Da as this is near the zero-crossing point for the NOE (28). To characterize their structures in solution, we therefore collected rotating frame data (ROESY) (28) as the signs of crosspeaks in this experiment are invariant to molecular size (**Fig. S1B**). We obtained roughly 30-50 distance constraints per peptide, or 10 NOEs per residue, for NMR structure calculations (**Table S2**). The resulting free peptide structures are relatively disordered extended conformations (**Fig. 1B**). However, we did observe some residual structure influenced by the side chain register. Specifically, tetrapeptides with a B-φ-B-φ motif (SS-31 and SPN4) had lower root-mean-squared deviation (RMSD) values (better structural precision) and three non-sequential NOEs, whereas those with a φ-B-φ-B motif (SS-20 and SPN10) lacked non-sequential NOEs (**Table S2**). Importantly, the non-sequential NOEs in SS-31 and SPN4 occurred between the two aromatic residues, consistent with previous observations that residual structure is more common in short peptides containing aromatic side chains (29, 30). Additional interactions that appear to stabilize the tetrapeptide structures in solution are cation-π interactions between neighboring basic and aromatic amino acids. These are supported by significant upfield ring current shifts for some of the basic residue side chain nuclei (**Table S1**), for all peptides except SS-31.

As a second approach to assessing peptide solution structures, we performed all-atom MD simulations (200 ns each) of the peptides in an aqueous environment. To initiate each simulation, the solution structure of each peptide closest to the mean of the NMR ensemble was used. To directly compare the ensemble of structures from our NMR- and MD-based approaches, we calculated the RMSD to the ensemble average and the radius of gyration (*R*_g_) values. For all peptides, the MD-derived ensemble had greater structural variability (higher RMSD) and were less compact (higher *R*_g_) than their cognate NMR-derived structures (**Fig. 1C**, compare black squares and circles). However, consistent with our NMR results, the RMSD values calculated by MD simulations were lower for SS-31 and SPN4 than for SS-20 and SPN10 (**Fig. S2**). Notably, all of these RMSDs were lower than the limiting RMSDs for tetrapeptides with random structures, which we calculate to be on the order of 3 to 5 Å (see Methods). Taken together, the results of our NMR and MD analyses suggest that the free peptides are largely disordered but retain some residual structure due to cation-π and aromatic ring stacking interactions. We next proceeded to empirically evaluate the interaction of our tetrapeptides with biomimetic membranes.

### Tetrapeptide analogs have distinct equilibrium binding behavior to cardiolipin-containing membranes

To assess the membrane binding properties of the peptide analogs, we performed isothermal titration calorimetry (ITC) of peptides titrated with large unilamellar vesicles (LUVs) containing a 20:80 molar ratio of 16:0/18:1 phosphatidylcholine (POPC) and 18:1 CL (tetraoleoyl-CL, TOCL) (**Fig. 2**). This approach provides a full thermodynamic characterization of the peptide-membrane interaction. Fits to binding isotherms (**Fig. 2A**) provided equilibrium binding parameters (**Fig. 2B**) that revealed key similarities and differences among the membrane-interactive properties of our test set. First, all tetrapeptides bound CL-containing membranes with roughly similar binding affinity (*K*_D_ 27.5 μM to 39.5 μM; Δ*G* −26.2 kJ/mol to −25.9 kJ/mol) that did not differ significantly among the analogs tested. However, the lipid-to-peptide binding stoichiometry, *n*, did differ among peptides. Compared with the benchmark SS-31 (*n* = 5.4; i.e., an average of 5.4 lipids per bound peptide), SPN4 had a similar value (*n* = 5.7), SS-20 had a significantly higher value (*n* = 7.4), and SPN10 had a significantly lower value (*n* = 3.3). These results indicate that SPN10 and SS-20 bind membranes, respectively, at higher and lower surface densities than SS-31.

**Figure 2.**
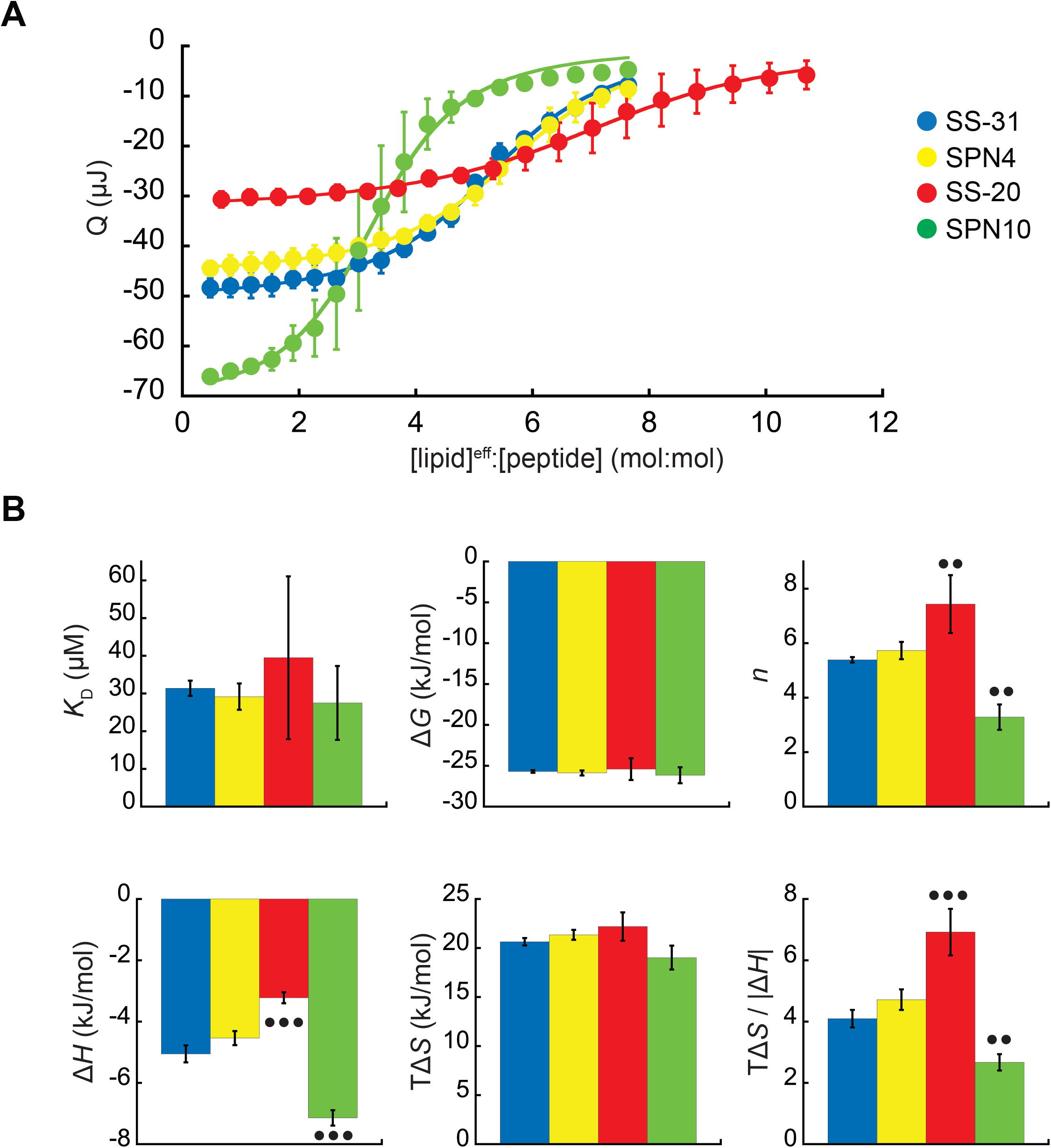
Microcalorimetry of peptide binding to LUVs. **A) Equilibrium binding isotherms.** Wiseman plots showing blank-corrected average integrated heats for lipid-into-peptide titrations as a function of [lipid]^eff^:[peptide] molar ratio using LUVs composed of an 80:20 molar ratio of POPC:TOCL with peptides color-coded as shown. Points represent means (n ≥ 3 ± SD) and curve fits are to binding models of single, independent sites. **B) Comparison of binding parameters.** Values of equilibrium binding parameters calculated from curve fits are shown for peptides color-coded as in Panel A. Statistical comparisons represent one-way ANOVA with Tukey’s multiple comparison test (α=0.05), with differences representing a comparison to SS-31 (no symbol, *P* > 0.05; • *P* ≤ 0.05; •• *P* ≤ 0.01; ••• *P* ≤ 0.001).

Membrane interaction of all peptides was enthalpically favorable (Δ*H*<0) but dominated by favorable entropy (TΔ*S*>0), as observed previously for SS-31 (18). In comparison with the binding enthalpy of SS-31 (Δ*H* = −5.1 kJ/mol), SPN4 had a similar value (Δ*H* = −4.5 kJ/mol), whereas the magnitude of binding enthalpy was significantly lower for SS-20 (Δ*H* = −3.2 kJ/mol) and higher for SPN10 (Δ*H* = −7.1 kJ/mol). As Δ*H* is a function of polar contacts made during binding (31), this trend in Δ*H* is consistent with the number of aromatic side chains containing polar groups (SPN10, two indole NH groups; SPN4/SS-31, one phenol OH group; SS-20, none). Binding entropy, which in this system is largely a function of aromatic side chain partitioning into the acyl chain region (31), did not differ significantly among peptides (TΔS ranged from 19.0 kJ/mol to 22.1 kJ/mol). However, the relative enthalpic and entropic contributions to binding, quantified by the ratio TΔS/|ΔH|, indicate that in comparison with SS-31, membrane binding of SS-20 is more entropy-driven and that of SPN10 is more enthalpy-driven. Having established these differences in membrane binding behavior among our peptide analogs, we then investigated their structural differences in the membrane-bound state.

### The bicelle-bound peptides adopt distinct reverse turn structures, except for SS-31

To determine the structural features of cationic-aromatic tetrapeptides in a membrane-like environment, we used bicelles as membrane mimetics. We hypothesized that moving from a high to a low dielectric environment would promote a more uniform structure and that the measured differences in the membrane-binding thermodynamics of these peptides could have a structural basis. As previously reported (4), in the presence of cardiolipin-containing bicelles, the NMR signals of SS-31 broadened at low peptide:lipid ratios, and then sharpened again at a molar excess of peptide to cardiolipin. This indicates that the free and bicelle-bound states of the peptides are in fast exchange on the NMR time-scale and should be amenable to transferred NOE (trNOE) studies (32). We confirmed this by observing that a large number of negative NOEs, consistent with a high MW peptide-bicelle complex, are transferred to the free peptide (**Fig. S1C, Fig. S3**). These trNOEs allowed us to calculate structures for all four peptides in their bicelle-bound states (**Table S3**), as shown in **Fig. 3**. The NMR structures are precise because they are each defined by 95-110 trNOEs per tetrapeptide, or about 25 structural restraints per residue (**Table S3**). This is reflected in heavy atom RMSDs values of 0.5-0.9 Å for the bound peptides, which is less than half of those of the free peptides (**Fig. 1C, Table S4**). NMR structures of all the peptide analogs had lower RMSD and *R*_g_ values in the membrane-bound relative to the free state. In other words, they became more structurally constrained and compact upon binding (**Fig. 1C**, compare black and red squares). The exception was SS-31, whose RMSD and *R*_g_ values did not statistically change upon binding.

**Figure 3.**
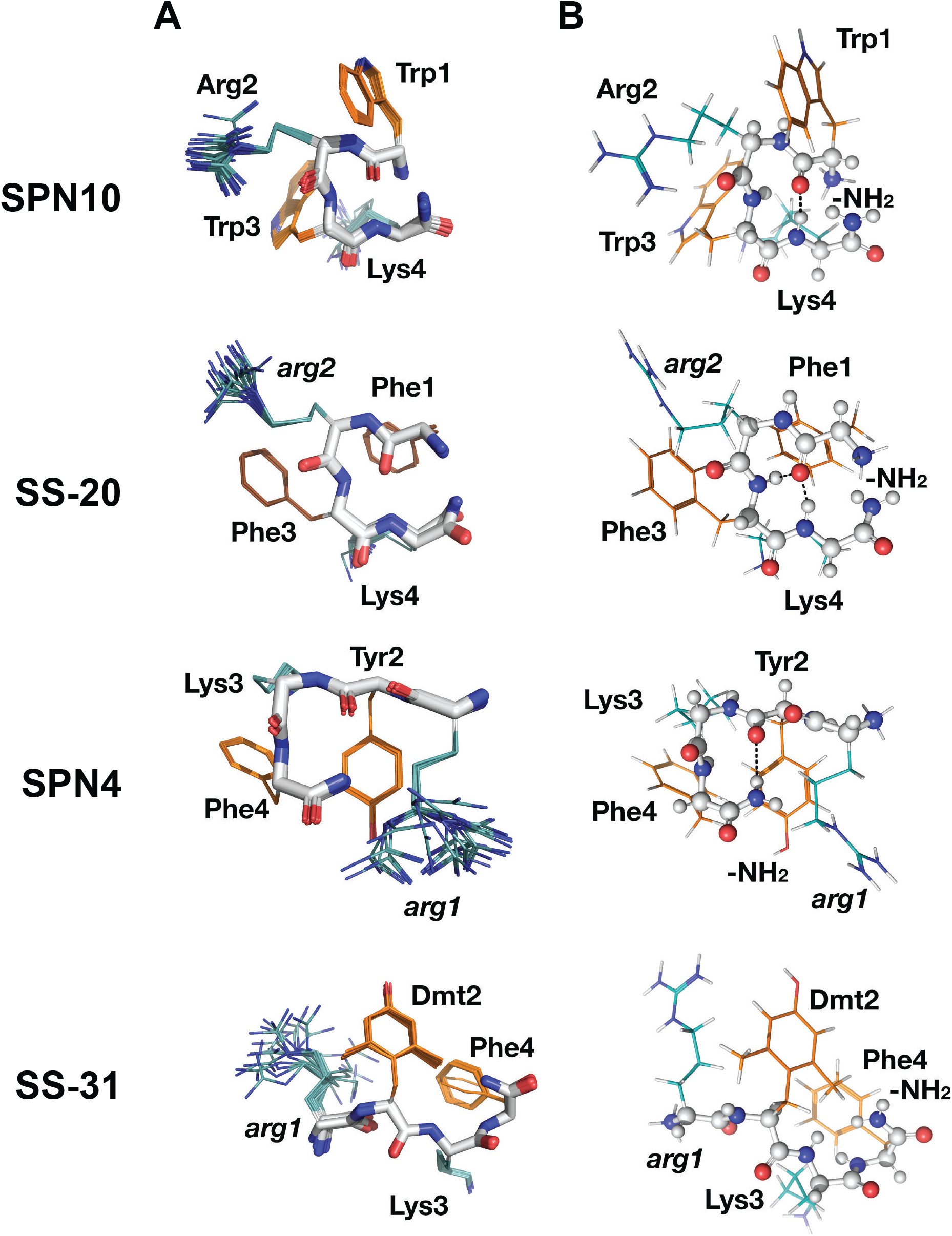
Peptide structures in the bicelle-bound state. **A) Top 20 lowest energy NMR conformers.** Backbone atoms and side chains are color coded as in Fig 1B. **B) Peptide secondary structure.** Conformations closest to NMR ensemble means are shown. Backbones are shown in ball-and-stick representation, H-bonds are shown as dotted lines, and side chains are shown as lines (orange for aromatic, teal for basic). Three of the peptides have main chain H-bonds: SPN10 [CO(1) to NH(4)], SS-20 [CO(1) to NH(4)], and SPN4 [CO(2) to NH_2_(5)]. SS-31 is extended and has no H-bonds.

Interestingly, we found that in the membrane-bound state, all peptide analogs except for SS-31 formed H-bonded reverse turn structures with basic side chains pointing away from the plane of the backbone ring (**Fig. 3A**). This gives the peptides a markedly asymmetric charge distribution (**Fig. S4**), with the cationic face of the peptides likely poised for binding to the negatively charged lipid phosphates of CL-containing membranes. To form the reverse turn structures, the φ-B-φ-B peptides have CO(1) to NH(4) H-bonds. However, for the B-φ-B-φ peptide SPN4, the H-bond is formed with the capping NH_2_ group that essentially acts as the amide proton donor of a non-existent fifth residue in a CO(2) to NH2(5) pattern (**Fig. 3B**). In contrast, the other B-φ-B-φ peptide SS-31 adopts an extended conformation due to steric restraints induced by the methyl groups on Dmt2 that preclude the turn conformation from forming. For the peptides that form a reverse turn, intra-peptide cation-π interactions are observed and the backbone H-bond NH donor in the turn structure is always an aromatic residue. For the φ-B-φ-B peptides SS-20 and SPN10, cation-π interactions form between Arg2, Phe3 and Trp3, Lys4, respectively. Although these cation-π interactions may arise simply from sequence proximity, they could play a critical role in reducing the overall polar character of these peptides, allowing them to traverse low-dielectric cellular structures. As for the basic residues, Arg is always the more poorly defined side chain in the structures and Lys is generally precisely defined, suggesting that the latter might be experiencing restricted motion due to partial insertion in the membrane (**Fig. 3A**).

We also evaluated the membrane-bound structures of the four tetrapeptides using MD simulations, where multiple peptides were allowed to associate with a bilayer consisting of an 80:20 molar ratio of POPC:TOCL (**Fig. 4A**). All peptides rapidly adsorbed to the membrane surface within 500-750 ns and evolved towards a stable, bound configuration over the course of the 2 μs simulations (**Fig. S5-S9**). As with the solution structures, membrane-bound structures from MD had higher RMSD and *R*_g_ values than those from NMR (**Fig. 1C, Table S4**).

**Figure 4.**
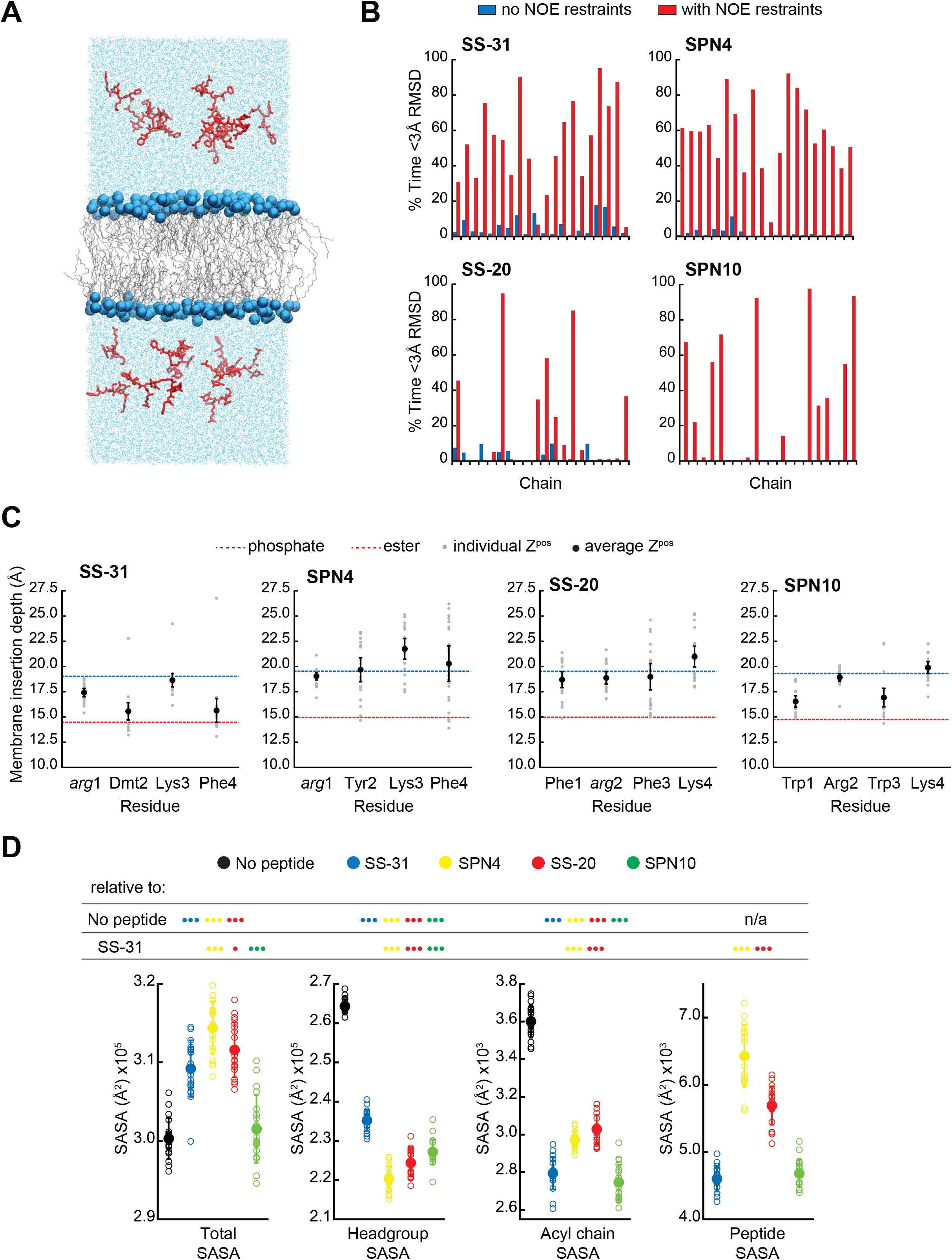
MD simulations of peptide conformations and membrane interactions. **A) Snapshot of a typical MD simulation.** Peptides (red) are shown in the aqueous phase on either side of a bilayer composed of an 80:20 molar ratio of POPC:TOCL (lipid acyl chains in wireframe, and lipid phosphates shown as blue van der Waals spheres. **B) Comparison of NMR and MD structures.** Comparison of the simulation time spent by each peptide analog below a heavy atom RMSD of 3Å to their respective lead NMR membrane-bound structures before (*blue*) and after (*red*) NOE restraints were imposed. **C) Average membrane insertion depths.** Bilayer depth (Z^pos^, n=20 ± 95% CI) for C^β^ atoms on each residue for each peptide shown in comparison with the average Z^pos^ levels of lipid headgroup phosphates (*dashed blue*) and lipid ester carbons (*dashed red*). The positions of the lipid atoms were averaged between the four different peptide systems for consistent comparison. The depths represent normalized distances to the bilayer COM. **D) Peptide-dependent SASA during MD trajectories.** SASA of the total bilayer and peptides system (*Total*) and individual components (*Headgroup*, *Acyl chain*, and *Peptide*) calculated from MD trajectories in the absence of peptide or with the peptide analogs indicated. Open symbols show individual block-averaged datapoints from each trajectory; solid symbols and error bars show means and SD for each dataset. Statistical comparisons are based on the Wilcoxon rank sum test, with differences representing a comparison to the No-peptide control, or to SS-31 as indicated (no symbol, *P* > 0.05; • *P* ≤ 0.05; •• *P* ≤ 0.01; ••• *P* ≤ 0.001).

To further compare our membrane-bound MD and NMR structures, we calculated the average fraction of time each peptide in the simulation was within 3Å heavy atom RMSD to the NMR top structure. Overall, the MD ensembles had a low frequency of sampling states with high similarity to the NMR-derived structures (**Fig. 4B, Table S5**). This low degree of structural overlap motivated the extension of each simulation for an additional 1 μs, during which NOE distance restraints between residues *i to i*+*2* and *i to i*+*3* were imposed for each peptide (33–35). Imposing the NMR restraints yielded considerable improvements in structural similarity to the NMR structure for all peptides (**Fig. 4B, Table S5**). NOE violations were computed over the MD simulations using time and ensemble averaging and violations occurred less than 20% of the time for the majority (∼71%) of the applied restraints (**Fig. S10**). Thus, all subsequent MD analyses were performed using NOE restrained MD data, unless otherwise indicated.

### Peptide analogs have different membrane insertion profiles and lipid interactions

We next analyzed our MD simulations for the membrane insertion depth of each peptide side chain. Side chain insertion depth (Z^pos^) for each peptide analog was measured by averaging the distance in the Z-direction (bilayer normal) from each side chain’s C*β* atom to the bilayer center of mass (COM) (**Fig. 4C, Fig. S5-S9**). Generally, the residues of SS-31 tended to bury deepest, followed sequentially by SPN10, SS-20, and SPN4.

In the trNOE experiments, we also observed NOEs between the peptides and the bicelle lipids, for which we used lipid proton assignments from the literature (**Fig. S11-S12**) (36). Most of the peptide-lipid NOEs were from aromatic residues to lipid protons close to the headgroup region. This suggests peptides are superficially buried in the interfacial region of lipid bilayers, in agreement with the MD results. Interestingly, the peptide-lipid NOEs for SPN10 were generally weaker than those of the other peptides (**Fig. S12**). Although we saw few, if any, trNOEs from lipid to the basic groups, this could be explained by the fact that the TOCL ^1^H signals were largely overlapped and dominated by the ∼40-fold molar excess of DHPC and POPC (**Fig. S11**). Additionally, given the 1/r distance dependence of charge-charge interactions, long-range electrostatic effects could be missed by the ∼5Å detection threshold of the NOE.

### SPN10 minimizes membrane surface area and CL self-interactions

We next used MD simulations to obtain insights into the effects of peptide binding on bilayer properties. We first considered molecular exposure at the membrane-solvent interface by calculating the solvent accessible surface area (SASA) of different groups (**Fig. 4D, Fig. S13**). We measured the total membrane and peptide SASA of a given system in addition to its component headgroup, acyl chain, and peptide SASA values in the presence and absence of peptides. In the absence of peptides, total membrane SASA can be parsed into headgroups, which constitute the majority of solvent-exposed area, and acyl chains, whose exposure can be interpreted as interfacial lipid packing defects (18, 37). As expected, the presence of all peptides at the bilayer interface reduced both lipid headgroup and acyl chain SASA, due to peptides “covering” the bilayer surface. Among the four analogs, SS-20 and SPN4 were more highly solvent exposed, consistent with their more superficial binding near the solvent interface (**Fig. 4C**). By comparison, SS-31 and SPN10 caused the greatest decrease in acyl chain SASA, consistent with their deeper burial into the nonpolar core (**Fig. 4C**), filling the “voids” of solvent-exposed hydrocarbon chains. The most notable distinction between the more deeply buried SS-31 and SPN10 was that SPN10 also caused a significantly lower headgroup SASA. Hence, SPN10-containing bilayers had the lowest total SASA among all peptide analogs, and SPN10 was the only peptide to not significantly change the total SASA compared to the bilayer-only system. The observation that total SASA is unchanged when SPN10 binds the bilayer indicates that the total amount of surface that is “created” by this bound peptide is approximately equal to the total amount of bilayer surface area that is “buried” by it.

Second, we considered peptide-induced elastic deformations of the bilayer, namely cross-sectional area per lipid and bilayer thickness (**Fig S14**). Our measurements of these parameters before peptide binding (i.e., peptides restrained in solution) were consistent with previous MD work from our group (18, 37, 38) and others (39). Upon peptide binding, we observed an inverse trend between bilayer thickness (**Fig. S14 A,B**) and mean lipid area (**Fig. S14 C,D**). SS-31 and SPN10 expanded mean lipid area (decreased bilayer thickness) to a greater extent than did SPN4 and SS-20, which is likely related to the deeper burial of SS-31 and SPN10 aromatic side chains into the membrane (**Fig. 4C**). The combined observations that SPN10 both expanded membrane area (**Fig. S14**) and maintained the lowest total SASA (**Fig. 4D**) among our peptide set suggests that even though this analog causes elastic bilayer expansion, it maintains a low “ruggedness” of exposed membrane surface (40).

Finally, we constructed radial distribution profiles to determine peptide-lipid and lipid-lipid interactions in the lateral (x-y) plane of the membrane (**Fig. S15**). As a proxy for peptide-lipid association (**Fig. S15**, *upper panels*), we chose the C*ζ* atom of the Arg at the first (SS-31/SPN4) or second (SS-20/SPN10) position. We observed three radial shells of POPC and TOCL phosphates around each Arg. SS-31 and SPN4 had a greater density of CL phosphates in the closest shells, whereas CL phosphates of distal radial shells were more populated for SS-20 and SPN10. This may be related to the proximity of the Arg to the cationic NH_3_ terminus in the B-φ-B-φ peptides causing greater local density of anionic CL. The distribution profiles of Arg-PC were, by comparison, much more similar among peptides. Our analysis of lipid-lipid radial distributions (**Fig. S15**, *lower panels*) revealed four radial densities of lipid phosphates around corresponding phosphates of CL or PC. CL phosphates in the same Z-plane do not often approach closer than ∼5 Å (18), likely due to charge-charge repulsion. Notably, SS-31 caused an increased density of CL around itself at distances < 6Å, suggesting that SS-31 may draw CL phosphates out of their respective Z-plane more than other analogs, perhaps consistent with this peptide’s membrane-thinning effects (**Fig. S15**). By comparison, SPN10 disfavored close contact of CL, which may be related to its large aromatic Trp side chains inhibiting close approach of sterically bulky CL.

### SPN10 has a markedly greater effect on membrane electrostatics

Our previous work showed that SS-31 modulates membrane surface electrostatics as a key part of its molecular MoA (18). We therefore sought to evaluate the effects of our four peptide analogs on membrane electrostatic potentials (**Fig. 5**). We first analyzed surface potential (Ψ_s_), which originates from fixed charges at the interface and is strongly negative for CL-rich mitochondrial membranes (41, 42). To this end, we used the fluorescent reporter probe ANS, which reversibly binds anionic membranes, with corresponding increase in quantum yield, in a manner that is promoted by Ψ_s_ attenuation (42, 43). As we have shown, ANS profiles are consistent with zeta potential readouts of membrane surface charge (18). We first measured the effect of peptide analogs on the surface charge of mitochondrial membranes by titrating mitoplasts (mitochondria with disrupted outer membrane) with peptide (**Fig. 5A**). All peptides caused a saturable decrease in membrane surface charge, with SPN10 causing markedly higher attenuation (**Fig. 5A**, *left*); by comparison, within the resolution of this assay, there was no discernible difference in saturation binding among peptides (**Fig. 5A**, *right*). These results support that the highest Ψ_s_ down-regulation is caused by SPN10 *in organello*, but do not rule out that this could be caused by peptide interaction with mitochondrial proteins. We therefore repeated this analysis in a more reductionist system with CL-containing LUVs (**Fig. 5B**). Again, SPN10 showed the greatest effect on Ψ_s_ in this lipid-only system (**Fig. 5B, *left*) despite all analogs having similar saturation curves (Fig. 5B**, *right*). Taken together, these results show that SPN10 is a markedly more potent attenuator of membrane surface charge, originating from down-tuning of lipid bilayer surface charge.

**Figure 5.**
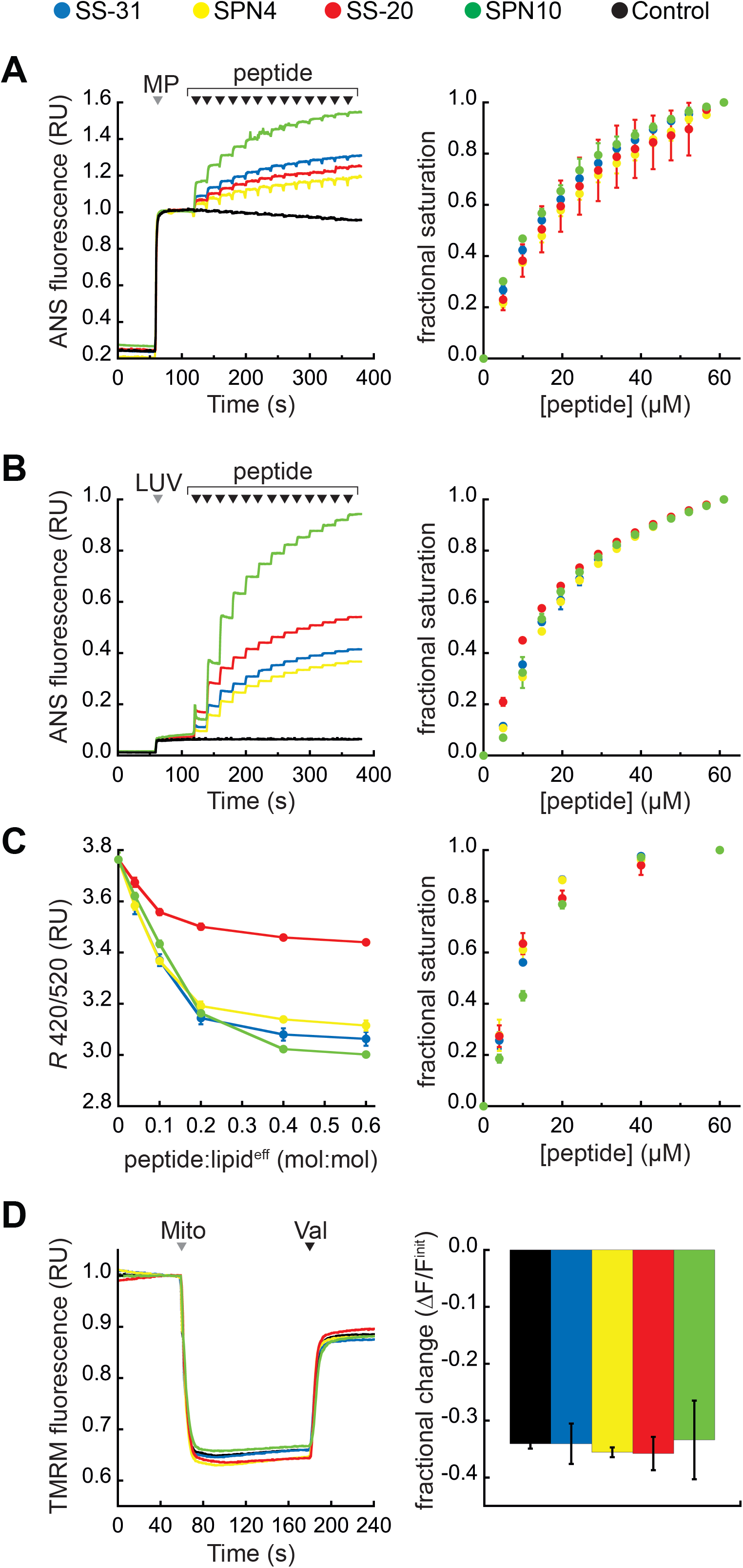
Effects of peptides on membrane electrostatic potentials. **A,B) Effects on surface potential (Ψ_s_).** *Left*, signal-averaged time courses of 1,8-ANS emission with addition of (A) mitoplasts (generated from *Saccharomyces cerevisiae* mitochondria, 200 μg total protein) or (B) 80:20 POPC:TOCL LUVs (100 nmol lipid^eff^), shown by *gray arrowheads* (“MP” and “LUV”, respectively), followed by sequential addition of peptide (10 nmol each), shown by *black arrowheads*. *Right*, saturation binding curves taken from 1,8-ANS time course data. **C) Effects on dipole potential (Ψ_d_).** *Left*, profile of di-8-ANEPPS measurements (*R*, ratio of 670 nm emission from 420 nm and 520 nm excitation peaks) in the presence of 80:20 POPC:TOCL LUVs (25 nmol lipid^eff^). *Right*, saturation binding curves taken from ratiometric di-8-ANEPPS measurements. **D) Effects on transmembrane potential (ΔΨ_m_).** *Left*, signal-averaged time course profiles of TMRM emission with addition of mitochondria from *S*. *cerevisiae* (200 μg total protein) following preincubation with 10 μM of respective peptide (*gray arrowhead*, “Mito”) and addition of ionophore valinomycin (*black arrowhead*, “Val”). *Right*, fractional change in TMRM emission following mitochondria addition. All means and traces are from n=3 independent samples and all error bars indicate SD. Control, addition of peptide buffer vehicle only.

We next analyzed membrane dipole potential (Ψ_d_), which originates from the arrangement of interfacial lipid and water dipoles, and contributes significantly (several hundred mV) to membrane electrostatic profiles (44) (**Fig. 5C**). Importantly, Ψ_d_ influences the translocation of hydrophobic ions across bilayers and may affect binding interactions with peptides (45). To evaluate the effects of our peptide analogs on Ψ_d_, we used ratiometric fluorescence excitation measurements of the membrane-bound probe di-8-ANEPPS, which has been shown to report dynamic changes in Ψ_d_, of model membranes (46, 47). Titration of LUVs with peptides resulted in a saturable reduction in Ψ_d_ (**Fig 5C***, left*) with fractional binding (**Fig 5C***, right*) consistent with our ITC-based binding curves (**Fig. 2**). These results indicate that all peptides cause saturable disordering of lipid and/or water dipoles upon binding. Furthermore, SS-20 had a markedly weaker effect than the others, possibly due to its comparatively lower aromatic bulk and/or lack of polar groups on aromatic side chains.

Lastly, we tested the effect of our peptides on the transmembrane potential (ΔΨ_m_), which is based on ion asymmetry across the IMM established by OXPHOS proton pumping (48). Both SS-31 and SS-20 have been shown to have no effect on ΔΨ_m_ in healthy mitochondria (3, 18), and it is crucial to verify that other mitochondria-targeted peptides have no IMM-uncoupling properties. To test this, we used the potentiometric probe TMRM, whose fluorescence quenches upon accumulation in the matrix of energized mitochondria (49). No peptides had any measurable effect on the TMRM-detected magnitude of ΔΨ_m_ (**Fig. 5D**), indicating that they neither hyperpolarize nor depolarize the IMM.

Taken together, these mitochondria-targeted tetrapeptides modulate membrane electrostatic potentials differently, in ways that depend on their side chain compositions. Most notable was the effect of SPN10 on Ψ_s_, which we propose to be a key underpinning of the activity of these peptides (18). Having addressed key aspects of their molecular structure and behavior, we proceeded to test how these features of our tested peptides relate to their efficacy in ameliorating stress using cell culture models.

### Cellular activity of CL-binding peptides

Therapeutic efficacy requires cationic-aromatic peptides to be cell-permeable. Although this has been extensively demonstrated for SS-31 (B-φ-B-φ) (2, 3), it was unclear if analogs with φ-B-φ-B sequence would behave similarly. We therefore compared the cell uptake and mitochondrial localization of N-biotinylated SS-31 and N-biotinylated SPN10, in human kidney epithelial cells (HK-2) and retinal pigment epithelial cells (ARPE-19). The two biotinylated peptides readily penetrated both cell lines within an hour and showed a mitochondrial localization pattern (**Fig. 6A**).

**Figure 6.**
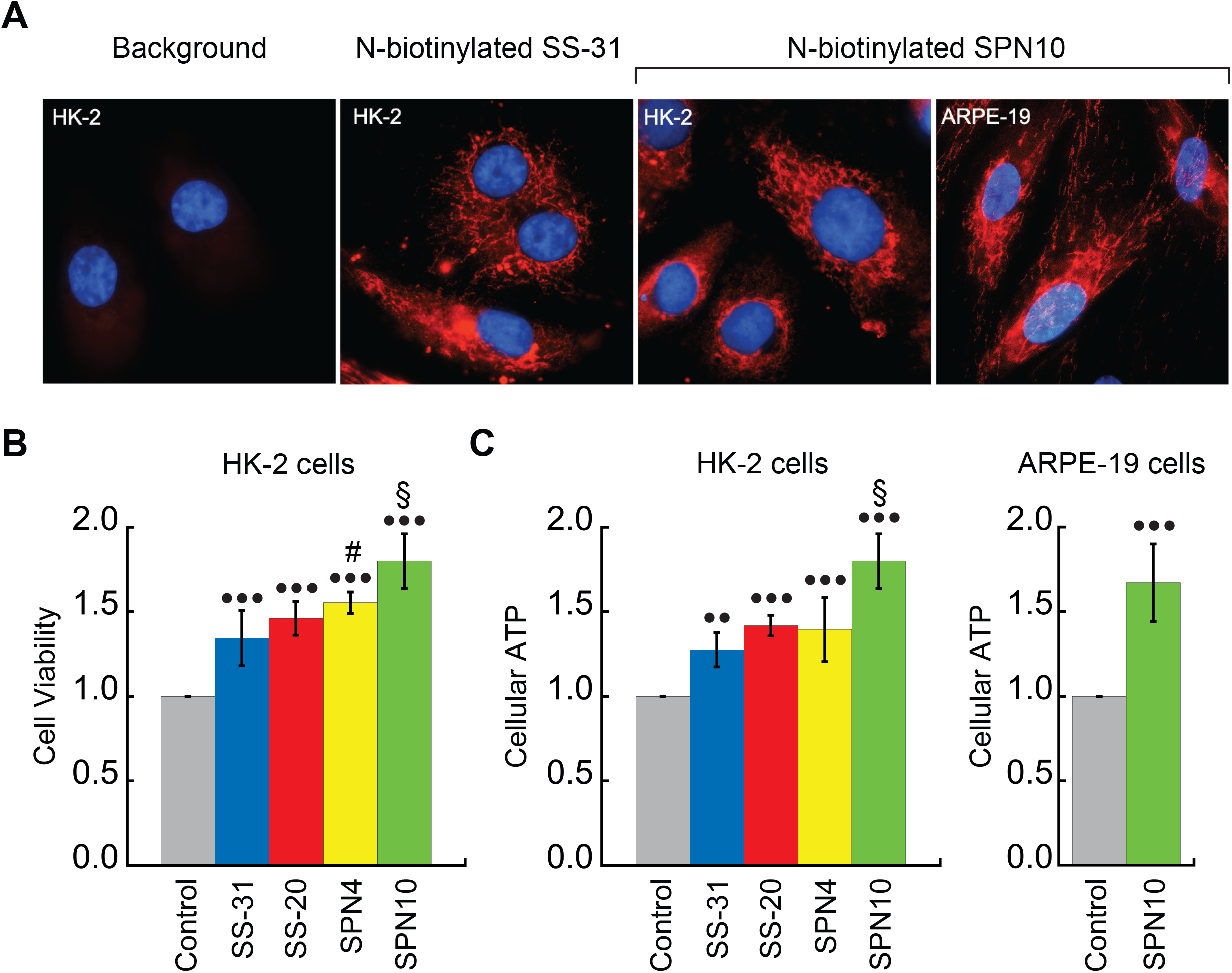
Cellular localization and efficacy of mitochondria-targeted peptides in cell culture. **A) Intracellular localization of peptides.** Confocal microscopy images of N-biotinylated variants of SS-31 and SPN10 in HK-2 and ARPE-19 cells as indicated. Biotin was visualized with streptavidin-AlexaFluor 594 and cell nuclei were stained with Hoechst 33342. **B,C) Peptide-dependent restoration of viability and energy metabolism in cell culture stress models.** HK-2 and ARPE-19 cells, as indicated, subjected to 7d of serum deprivation in the absence or presence of mitochondria-targeted peptides (10 nM) showed peptide-dependent increase in B) cell viability, and in C) cellular ATP levels. Statistical comparisons represent one-way ANOVA with Tukey’s multiple comparison test, with differences representing a comparison to vehicle-only control (•• *P* ≤ 0.01; ••• *P* ≤ 0.0001) or comparison to SS-31 treatment (# *P* ≤ 0.05; § *P* ≤ 0.001).

Extensive studies have shown that SS-31 promotes cell survival under a variety of stress challenges, including ischemic, hypoxic, metabolic, and oxidative stress conditions (10). Serum starvation is frequently used as a stress model in cell cultures, since it causes decreases in cellular ATP, cell cycle arrest, and apoptosis (50–52). The effects of the four test peptides on cell viability and ATP levels were examined in HK-2 cells seven days after serum removal. Treatment with the four peptide analogs at concentrations of 10 nM significantly increased cell viability in HK-2 cells, with SPN10 being significantly better than SS-31 (P < 0.001) (**Fig. 6B**). Cellular ATP content was also significantly elevated by the four peptides, and SPN10 was again better than SS-31 (P < 0.001) (**Fig. 6C**). In addition, SPN10 significantly raised ATP levels in ARPE-19 cells with serum starvation (**Fig. 6C**).

## Discussion

The objective of this study was to evaluate the effects of modifying two key features of mitochondria-targeted peptides, side chain composition and sequence register, to better understand what chemical properties are important for their therapeutic efficacy. SS-31 and SS-20, which have both been validated in pre-clinical and clinical tests, feature different aromatic side chains and ordering of cationic and aromatic groups. The efficacy of these compositionally distinct peptides motivated us to test the structural and functional consequences of further modifications in the cationic-aromatic motif of these peptides. All four analogs were found to bind CL-containing bilayers, modulate membrane electrostatics, and show efficacy in cell stress models. Collectively, these observations support the concept that this general motif is sufficient for the molecular action of these mitochondria-targeted compounds. Yet there were also marked differences in their structures and molecular behaviors. These differences provide critical insights into the MoA of this class of compounds that can be leveraged to develop more effective mitochondrial therapeutics.

A central aim of this study was the structural characterization of these peptides in solution and in the membrane-bound state. To this end, we used complementary approaches of NMR spectroscopy and MD simulations, providing the first models of the structures of these compounds. As expected, all peptides are largely disordered in solution (**Fig. 1B**, **Fig. S2**) and assumed a more compact and well-defined structure when membrane-bound (**Fig. 3**). This trend is reflected by the general decrease in the calculated RMSD and R*_g_* seen for the bound structures (**Fig. 1C**). Intra-peptide aromatic ring stacking and cation-π interactions are stabilizing features of both the solution and membrane-bound states (**Fig. 1B**, **Fig. 3**). This point is significant because cation-π complexes are known to lower the energetic cost of partitioning basic side chains into the nonpolar membrane core (53, 54), which likely contributes to the ability of these tetrapeptide analogs to traverse cell membranes, including those of the blood-brain barrier (2, 55), and to reside stably at the membrane interface.

The most notable structural feature of these tetrapeptides is the membrane-bound reverse-turn conformation observed for all analogs other than SS-31. These conformers can all be formally classified as four-residue beta turns stabilized by an *i* to *i*+3 main chain H-bond with an Cα(1) to Cα(4) distance of ∼7Å or less (56, 57). The dihedral angles of the *i*+1 and *i*+2 residues of these structures deviated from the canonical ϕ/Ψ values for the most common beta turns; however, with increasing availability of high-resolution protein structures, it is becoming clear that the central residues of beta turns can occupy a wider diversity of Ramachandran space than previously appreciated (58). The φ-B-φ-B peptides (SS-20 and SPN10) are stabilized by a CO(1) to NH(4) H-bond, whereas the B-φ-B-φ peptide (SPN4) is stabilized by an H-bond between CO(2) and the C-terminal amide group (in lieu of a ‘fifth’ residue) (**Fig. 3**). This latter feature underscores the importance of C-terminal amidation in these compounds: not only does the amide remove the C-terminal carboxyl function to maintain the net +3 charge of these compounds, but as shown in this study, it also provides an H-bond donor to stabilize the turn structure of peptides with B-φ-B-φ register. Given that the H-bonds of these reverse turn structures reside in the low-dielectric microenvironment of the membrane interface, they are likely to be more stabilizing than when in bulk aqueous solution (59). By partially satisfying the H-bonding capacity of main chain atoms, this *i* to *i*+3 polar interaction partially offsets the energetic penalty of dehydration of polar backbone functional groups at the membrane, which may in turn allow the peptides to reside near the polar-apolar boundary of the membrane. From these findings, one possibility to explore is the synthesis of cyclized forms of these peptides, given that forcing the peptides to adopt the bioactive pose may lessen the entropic penalty of membrane binding.

Our results provide several insights into the nature of the interaction between the tetrapeptides and lipid bilayers. The first insight pertains to peptide binding density (**Fig. S16**). Based on the lipid:peptide stoichiometries (*n*) from our ITC analyses (**Fig. 2**), coupled with the known cross-sectional areas of POPC (70 Å^2^, (60)) and TOCL (129 Å^2^, (37)), one can calculate the binding footprint of peptides in the bilayers (80:20 POPC:TOCL) used for ITC. In increasing order, the per-peptide membrane areas are: 270 Å^2^ (SPN10) < 434 Å^2^ (SS-31) < 466 Å^2^ (SPN4) < 605 Å^2^ (SS-20). Hence, SPN10 has greater binding density, and SS-20 has lower binding density, compared with SS-31/SPN4. Given that the mitochondrial IMM is protein-rich with little free exposed bilayer (61), the greater per-peptide membrane coverage of SPN10 may relate to its enhanced efficacy by increased occupancy of the limited lipid area of mitochondria (**Fig. S16**). This leads to a second related insight from this work regarding the effects of these peptides on membrane surface area. Our previous work showed that SS-31 caused a decrease in acyl chain SASA, likely related to a decrease of interfacial hydration and increased lipid packing density (18). In the present study, we found that SPN10 stood apart from the other peptides in that it minimized solvent exposure of both lipid aliphatic chains and headgroups, thereby resulting in total membrane SASA similar to membranes in the absence of peptide (**Fig. 4D**). This ability of SPN10 to fill packing defects is likely related to a combination of factors, including its large aromatic side chain volume, its deeper binding in the membrane interface (**Fig. 4C**), and its high surface coverage (**Fig. 2**). Within the IMM, the inverted conical geometry of CL creates lateral packing defects of lamellar bilayers (37), which could be related to transient pore-like defects that allow small molecule permeation (62) and/or accessibility of acyl groups to pro-oxidants that cause lipid peroxidation (63). Reducing lipid packing voids could be part of the MoA of these tetrapeptides, contributing to the observed decrease in proton leak across the IMM (11) and lipid oxidative damage (1, 64–66) that occurs with SS-31 treatment. A final related point pertains to specific peptide-lipid interactions. Our trNOE (**Fig. S12**) and MD (**Fig. 4C**) results both indicate that bound peptides reside in the membrane interface, likely within the boundaries of the lipid phosphate and ester groups. Yet the peptide analogs may mediate different lipid interactions. Compared with other peptides, the NOEs of SPN10 to lipid protons were notably fewer and weaker (**Fig. S12**). This may be related to our MD radial distribution profiles showing that the Arg of SPN10 has a preferential lateral accumulation of POPC over TOCL (**Fig. S15**). Understanding exactly how specific side chain-lipid chemical interactions relate to the efficacy of these tetrapeptide analogs will require further investigation.

Our calorimetric binding analysis showed unexpected differences in the thermodynamics of the interactions of these peptides with CL-containing membranes (**Fig. 2**). All analogs had roughly equal *K*_D_ (ΔG) values, with binding dominated by favorable changes in entropy (mean TΔS/|ΔH| ranged from 2.7 to 6.9). However, compared with SS-31/SPN4, the binding of SPN10 and SS-20 had larger and smaller enthalpic binding components, respectively. These features may be interpreted in terms of the origins of ΔS and ΔH for peptide-membrane interactions. First, the entropic cost of small peptide binding that comes from restricting conformational, translational and rotational degrees of freedom (adsorption entropy) is likely to be small based on theoretical considerations (67). Instead, the large and favorable binding entropy we observe originates largely from the classical hydrophobic effect: membrane penetration of aromatic side chains is attendant with increased solvent mobility that accompanies the desolvation of the peptide and the release of ordered waters from nonpolar acyl surfaces (31). We propose that membrane binding of SS-20 is dominated by entropically favored burial of its aromatic Phe side chains, which lack polar groups, in the nonpolar membrane core. This may facilitate deeper burial of SS-20 aromatic rings in the acyl chain region, consistent with its larger binding footprint. By comparison, binding enthalpy originates largely from polar interactions between peptide and lipid headgroups. We propose that membrane binding of SPN10 is strongly favored by polar interactions involving the indole NH hydrogen bond donors of its Trp side chains. Trp is known to remain partially in the interfacial region due to its large rigid paddle-like structure (68–72), which may be consistent with its higher observed binding density. Finally, our analysis suggests that the interaction of these peptides with membranes shows enthalpy-entropy compensation, wherein the binding energy of a congeneric series of compounds remains relatively constant due to opposing changes in ΔH and ΔS (73), as observed with the surface interactions of different peptides (74, 75). Hence, from a molecular engineering perspective, optimizing the efficacy of these tetrapeptides might be directed less toward enhancing binding affinity *per se* and more toward enhancing the enthalpic contribution, particularly by modulating polar contacts among aromatic groups.

This work revealed striking differences among the peptides in terms of their effects on membrane electrostatic potentials (**Fig. 5**). First, the Ψ_s_ of negatively-charged membranes (biomimetic liposomes and mitoplasts) was attenuated by all peptides (**Fig. 5A,B**); however, SPN10 had the greatest effect by far. This may be related in part to the higher binding density of SPN10. However, it may also be related to SPN10 uniquely having two aromatic side chains with polar (indole NH) groups that can each mediate H-bond interactions with lipid phosphates, which can alter ionization behavior and reduce headgroup charge (76, 77). Second, the Ψ_d_ of bilayers, related to the ordering of lipid polar groups and interfacial waters, was down-regulated by all peptides (**Fig. 5C**). This may be due to a general effect of bound peptide causing disorganization of interfacial water dipoles. However, as a group, those peptides with polar groups on their aromatic side chains (SPN4, SS-31, SPN10) attenuated Ψ_d_ much more than the peptide lacking aromatic polar groups (SS-20); hence, it is possible that polar contacts with lipid mediated by aromatic side chains may alter the orientation of lipid headgroup dipoles (e.g., the P-N vector of the phosphate-choline dipole of PC (78)). Finally, it is notable that no peptide affected the ΔΨ_m_ (**Fig. 5D**), supporting that they do not depolarize mitochondrial membrane potentials. Taken together, this work supports our working model (18) that the tuning of Ψ_s_ is a key part of the MoA of these peptides, correlated with efficacy.

Our evaluation of these peptide analogs in cell culture studies provided the most relevant ranking of their relative effectiveness (**Fig. 6**). All analogs targeted mitochondria (**Fig. 6A**) and, to varying degrees, restored viability and ATP content in serum starvation models of cell stress relative to vehicle-only control (**Fig. 6B,C**). In these tests, analogs SPN4, SS-20 and SPN10 consistently outperformed SS-31; based on our structural analysis, this difference in efficacy could be related to the ability of the analogs to form compact folds when membrane-bound. Most notably, SPN10 showed the greatest potency in cell stress recovery.

In summary, as the first structure-activity analysis for this class of compounds, this study provides new insights to guide their potential optimization. Given the complexity of membrane interactions in the molecular MoA of these compounds (18), coupled with the fact that membrane protein interactions are involved in their activity (19), our limited test set of four analogs could not unequivocally address all chemical features that may enhance function. But insofar as the composition and activity of SPN10 could provide a direction for optimization, the engineering of future analogs may be guided by: (i) greater bulk of aromatic R groups (which, among proteinogenic amino acids, means emphasizing Trp); (ii) ability to form compact (reverse-turn) structures when membrane-bound; (iii) polar groups on aromatic side chains that enhance enthalpy of membrane interactions; (iv) ability to decrease SASA of lipid groups; and (v) ability to down-regulate membrane Ψ_s_. Together, modulation of these features will help pave the way for rational design of next-generation variants of this class of mitochondria-targeted therapeutic compounds.

## Materials and Methods

### Reagents

Peptides SS-31, SS-20, SPN4 and SPN10 were prepared by solid-phase synthesis as TFA salts by Phoenix Pharmaceuticals (Burlingame, CA). Powder stocks were reconstituted to a concentration of 10 mM as aqueous solutions and stored at −20°C. Synthetic phospholipids were purchased as chloroform stocks form Avanti Polar Lipids (Alabaster, AL), including POPC (1-palmitoleoyl-2-oleoyl-*sn*-glycero-3-phosphocholine), DHPC (1,2-diheptanoyl-*sn*-glycero-3-phosphocholine), and TOCL (1′,3′-bis[1,2-dioleoyl-*sn*-glycero-3-phospho]-*sn*-glycerol). All lipid stocks were stored at −20°C in clear glass vials with Teflon-lined cap closures. Fluorescent probes TMRM, ANS, and di-8-ANEPPS were purchased from Thermo Fisher Scientific (Waltham, MA). All solutions were prepared with ultrapure water (Millipore Advantage A10 system; resistivity 18.2 MΩ•cm @ 25°C; total oxidizable carbon ≤ 4 ppb).

### Isothermal Titration Calorimetry

ITC measurements were performed based on well established procedures (79) that we have previously used to measure peptide-membrane interactions (18). Solutions of peptide (titrate) and LUVs (titrant) were prepared in 20 mM HEPES-KOH, pH 7.5, and lipid-into-peptide titrations were performed with a low-volume nano-ITC microcalorimeter (TA Instruments, New Castle, DE). The calorimeter cell (volume 170 μl) contained 125-175 μM peptide and LUVs (20 mol% TOCL / 80 mol% POPC, 8 mM total lipid) were injected in aliquots of 2.5 μl (20 total injections) at time intervals of 300 s at 25 °C. To account for heats of dilution, experiments were performed by the addition of titrant into solutions of buffer only, which were used for baseline subtraction. Data from dilution-corrected and integrated heat flow time courses were fit as Wiseman plots (modeled as independent, identical single binding sites), from which equilibrium binding and thermodynamic parameters (*K_d_*, *n*, Δ*H*, and Δ*S*) were determined by nonlinear regression fits (NanoAnalyze software version 3.10.0, TA Instruments).

### Fluorescence Spectroscopy

Steady-state fluorescence measurements were performed with a Fluorolog 3-22 spectrofluorometer (HORIBA Jobin-Yvon, Edison, NJ) equipped with single photon-counting electronics, double-grating excitation and emission monochromators, automated Glan-Thompson polarizers, and a 450-watt Xenon short arc lamp. Measurements were made either in 4 × 4-mm quartz microcells or in 1 × 1-cm quartz cuvettes with a stir disc seated in a thermostated cell holder.

### Spectral Measurements of Membrane Electrostatic Potentials

Measurements of Ψ_s_, Ψ_d_, and ΔΨ_m_ were made with LUVs (80:20 POPC:TOCL), with active mitochondria isolated from *S*. *cerevisiae* as described (80), or with mitoplasts prepared by osmotic rupture (80), as indicated. Ψ_s_ measurements were performed as described using the 1,8-ANS reporter probe (18). Briefly, stirred reactions containing 0.95 μM 1,8-ANS (added from 10 mM methanol stock) and either LUVs (in 20 mM HEPES-KOH, pH 7.5 with [lipid]^eff^ = 50 μM) or mitoplasts (0.1 mg/ml total protein) were titrated with stepwise additions of 10 nmol peptide over 380 s time course measurements (λ_ex_ = 380 nm; λ_em_ = 460 nm). ΔΨ_m_ measurements were performed as described using the TMRM potentiometric probe (18). Briefly, stirred reactions of TMRM assay buffer (20 mM Tris-HCl, pH 7.5, 20 mM KCl, 3 mM MgCl_2_, 4 mM KH_2_PO_4_, 250 mM sucrose, 0.5% (w/v) fatty acid-free BSA), respiratory substrate (2 mM NADH), and 0.1 μM TMRM (added from 10 μM stock) were supplemented with mitochondria (0.1 μg/ml total protein) that were pre-incubated with or without 10 μM of peptide, followed by potential dissipation with 2.5 μM valinomycin over 240 s time course measurements (λ_ex_ = 546 nm; λ_em_ = 573 nm). Ψ_d_ measurements were performed as described using the di-8-ANEPPS reporter probe as described (46). Briefly, LUVs were prepared by adding 1 mol% di-8-ANEPPS (from ethanol stock) to phospholipids prior to drying lipid films under nitrogen gas, hydration and extrusion. Solutions with di-8-ANEPPS-containing LUVs (in 20 mM HEPES-KOH, pH 7.5 with [lipid]^eff^ = 100 μM) were titrated with peptide at the indicated peptide:lipid molar ratios and read by excitation scans (λ_ex_ = 380-580 nm; λ_em_ = 573 nm; 1 nm increments and 1 s integration times), from which the ratiometric value (*R*, emission resulting 420 nm: 520 nm excitation) was used as a readout of Ψ_d_.

### MD simulations

A similar approach was used as described previously (Mitchell et al., 2020) where the SS-31 peptide structure was generated by modifying an extended tetrapeptide with the sequence Arg-Tyr-Lys-Phe. Coordinates were modified using the VMD Molefacture Plugin (81) to invert the stereochemistry of the N-terminal Arg residue from L to D and to replace the 2′ and 6′ hydrogen atoms of the Tyr side chain with methyl groups. Parameters for the 2′,6′-Dmt were modeled after the parameters of 3′,5′-dimethylphenol after running its structure through ParamChem’s CGenFF server (82). Since our initial parameterization of SS-31 (18), CHARMM36m forcefield parameters for cation-π interactions have been developed. We have included cation-π terms for SS-31 and other peptide analogs to more accurately model interactions between aromatic side chains and choline head groups found in the bilayer interfacial region.

Tetrapeptides with amino acid sequences matching SS-20 (Phe-Arg-Phe-Lys), SPN4 (Arg-Tyr-Lys-Phe), and SPN10 (Trp-Arg-Trp-Lys) SS peptide analogs were initially generated with proper stereochemistry using the UCSF Chimera Build Structure Plugin (83). CHARMM-GUI was then used to amidate each peptide’s C-terminus, obtain CHARMM36m forcefield parameters with cation-π interactions enabled for aromatic residues (84–88), solvate each system with TIP3P water model and a 150 mM NaCl concentration, and generate simulation input files. Following the CHARMM-GUI standard protocol for solvated proteins, the peptide systems were energy-minimized using the steepest-descent algorithm for 5000 steps, followed by canonical (NVT) ensemble equilibration for 250 ps with a 1 fs timestep. All minimization, equilibration, and production simulations were performed using the GROMACS version 2019 (33, 34). These minimized and equilibrated structures of SS-20, SPN4, and SPN10 were used in our peptide-bilayer systems.

MD simulations were then used to characterize the binding process of the four peptide analogs and to investigate their respective effects on membrane structure and dynamics. All-atom systems with explicit membrane and solvent were prepared using CHARMM-GUI with the CHARMM-36m forcefield and the TIP3P water model (84–86, 88, 89). Bilayers were generated with TOCL and POPC lipids at a molar ratio of 20:80 TOCL: POPC. Each system contained a total of 150 lipids (75 per leaflet). Peptide-bilayer systems were constructed by removing solvent from the bilayer systems generated by CHARMM-GUI, resizing the box’s Z-dimension to 16 nm, placing 20 peptides (10 peptides on either side) 2-3 nm away from the bilayer’s headgroup region, and then re-solvating to ∼75% water by mass, and raising the salt concentration to 100 mM NaCl. Each system contained ∼93,000 atoms. A representation of the initial setup of the peptide-bilayer systems is shown in **Fig. 4A**. Following the CHARMM-GUI standard protocol for protein-bilayer systems, all systems were energy minimized for 5000 steepest descent steps, followed by canonical ensemble (NVT) equilibration for 100 ps with a 1 fs timestep, 200 ps of NPT equilibration with a 1 fs timestep, and ∼100 ns of NPT equilibration with a 2 fs timestep. The NPT equilibration steps were performed with semi-isotropic pressure coupling and the Berendsen barostat and the Berendsen thermostat. Position and dihedral restraints were used during equilibration on the lipids and peptides to maintain lipid geometry and bilayer morphology and to prevent the peptides from interacting with the bilayers during equilibration. To enforce an equal number of peptides interacting with each side of the bilayer (10 peptides per leaflet; 7.5:1 lipid-to-peptide ratio and a 1.5:1 cardiolipin-to-peptide ratio), two inverted flat-bottom restraints in the Z-direction were placed at the bottom of the box (Z=0 nm). A restraint was placed on the peptides’ Cα atoms with a force constant of 200 kJ mol^−1^ nm^−1^ and a radius of 3 nm, which served to maintain a constant lipid-to-peptide ratio on either side of the bilayer. A second restraint was placed on the POPC phosphates with a force constant of 50 kJ mol^−1^ nm^−1^ and a radius of 4.5 nm to prevent the upper and lower leaflet from drifting in the z-direction, while still allowing for natural membrane deformations.

Production simulations were run for 2 μs and saved every 1 ps. In all production simulations electrostatic and Lennard-Jones (LJ) interactions were cut off at 1.2 nm, with electrostatics shifted from 0 nm to the cutoff, and LJ interactions shifted from 1.0 nm to the cutoff. Long-range electrostatic interactions were computed using the particle mesh Ewald method and a fourier spacing of 0.12 nm. All bilayer system production runs were simulated in the NPT ensemble using the Nose-Hoover thermostat, Parrinello-Rahman barostat, and semi-isotropic coupling scheme, with the temperature maintained at 303.15 K and pressure kept at 1.0 bar with semi-isotropic coupling. The time constants for pressure and temperature coupling were 5.0 and 1.0 ps, respectively, and the compressibility value was 4.5 E-5 bar^−1^. Simulations were performed using periodic boundary conditions in all dimensions and the simulation time step was 2 fs.

Following the 2 μs unrestrained production simulations additional 1 μs simulations were performed using NMR-derived NOE distance-restraints. Only those NOE restraints between residues *i* to *i+2* and *i* to *i+3* NOEs were imposed in the MD simulations. By only including “long-range” restraints we aimed to enforce the long-distance NOEs while still allowing for natural dynamics and not over-restraining the ensemble. A force constant of 5000 kJ mol^−1^ nm^−1^ was applied equally to each peptide (to promote sampling of more conformations) and averaged over the ensemble of 20 peptides. Since distance restraints based on instantaneous distances can heavily reduce conformational dynamics, the 20 peptides were restrained to a time-averaged distance with a decay time of 10 ps. The initial configuration for the NMR-restrained simulations was the final configuration of the 2 μs unrestrained simulations. An example .mdp file for the 20-peptide system with NOE restraints is included in the supporting information.

Simulations of single peptides in solution were also performed. The initial peptide structures were taken from the top structure in the NMR ensemble obtained in this study. The systems were solvated with the TIP3P water model and 100 mM NaCl salt concentration. These solvated systems were energy-minimized using the steepest-descent algorithm for 50000 steps (or a maximum force tolerance of 1000 kJ mol^−1^ nm^−1^), followed by canonical ensemble equilibration for 100 ps with 2 fs timestep, and 100 ps of NPT equilibration with a 2 fs timestep accomplished using the Parrinello-Rahman pressure coupling scheme and the V-rescale thermostat (90). Production simulations for each analog were run for 200 ns with a 2 fs timestep in an NPT ensemble using the Parrinello-Rahman pressure coupling scheme and the V-rescale thermostat.

### Analysis of MD simulations

The binding time-dependence, membrane insertion depth, and bilayer thickness were analyzed using the MDTraj Python module (91) to process the trajectories and in-house scripts using the NumPy (92, 93), SciPy (94), and Pandas (95) Python modules to perform calculations.

Average structures (S_avg_) were calculated using the *gmx rmsf* function in GROMACS for each of the five conditions (**Fig. 1C**) according to the following: one S_avg_ from 20 lead structures (both NMR conditions), one S_avg_ for each of the 20 peptides (MD: Membrane, 1-2 μs time range; MD: Membrane (restrained) 2-3 μs), and one S_avg_ for 13 × 15 ns intervals of the trajectory (MD: Solution; block averaging, see **Fig. S17**) (33, 34). The RMSD of the heavy atoms to their respective S_avg_ was calculated for each condition’s corresponding ensemble of peptide structures. The radius of gyration (R_g_) was calculated using the *gmx gyrate* function in GROMACS for the ensembles in each condition described above (33, 34).

NOE violations were calculated using in-house Python scripts that accounted for ensemble and time averaging (33–35). The instantaneous ensemble averaged distance (*r*(t)*) was computed according to equation 1

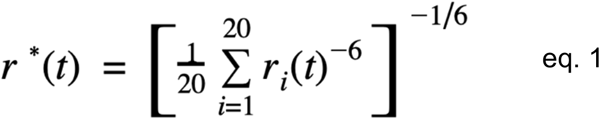

Time averaging was then performed according to equation 2

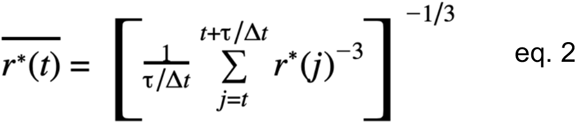

where *τ* = 10 ps, and Δt=1 ps. Violations were determined as those time- and ensemble- averaged pair distances which exceeded the experimentally determined upper bound NOE distances plus a buffer tolerance of 0.3 A. The list of NOEs implemented in the MD simulations and the fraction of time spent in violation of the upper bound for each NOE in each analog are presented in **Fig. S10**.

Solvent accessible surface area (SASA) measurements were calculated using the *gmx sasa* function in GROMACS (33, 34), which uses the double cubic lattice method (96). For the SASA analyses, we defined the acyl chain region of a lipid as the carbon and hydrogen atoms below the ester carbon. The area per lipid was derived from the *gmx energy* function in GROMACS (33, 34), which outputs the change in the X-Y box dimensions over time; the latter measurement was divided by the number of lipids in one leaflet (75 lipids). The RMSD of each simulated peptide analog in solution with reference to its initial structure was calculated using the *gmx rms* function in GROMACS (33, 34). Individual peptides were used as independent samples for calculating variance in the analysis of peptide-membrane MD simulations and the NMR RMSD and R_*g*_ measurements in **Fig. 1C**. Block averaging (97) was used to select decorrelated time intervals (**Fig. S17**) and calculate the standard deviation for all SASA measurements (**Fig. 4D**), bilayer thickness, area per lipid, and RMSD/R_*g*_ from MD simulations of a single peptide in solution. All images of the systems were created using VMD (81). All figures were created using the Matplotlib Python module (98).

### NMR sample preparation

For studies on the free peptides, samples were 10 mM in peptide at pH 6. For studies on the bicelle-bound peptides, bicelles were prepared according to a published method (4). The lipids DHPC, POPC, and TOCL were mixed at concentrations of 4.5, 1.5, and 0.15 μmoles, respectively, followed by drying under nitrogen gas for 20 minutes. After vacuum desiccation overnight, the dry lipid films were resuspended in 1 mL of de-ionized water, incubated at room temperature for 30 minutes, and gently swirled into solution. The peptides were added to the bicelles at a molar ratio of 5 peptide to 1 cardiolipin (TOCL), and the pH was adjusted to 5.5 with potassium hydroxide.

### NMR spectroscopy

NMR experiments were performed on a 600 MHz instrument equipped with a cryogenic probe. NMR assignments for the free peptides were obtained from 2D TOCSY (t_m_ = 70 ms), ROESY (t_m_ = 200 ms), and NOESY (t_m_ = 150 ms) spectra, where t_m_ is the mixing time. These were supplemented with ^13^C-HSQC and ^15^N-HSQC spectra obtained for the peptides at natural isotope abundance. All data were obtained at a sample temperature of 25°C Assignments for all four peptides are given in Table S1. Chemical shift deviations from random coil values (99) were used to infer cation-pi interactions. NMR structures of the free peptides were calculated using distance restraints obtained from ROESY spectra with t_m_ = 200 ms. For transferred NOE experiments, the free and bound peptides are in fast exchange and trNOE experiments were performed on samples that had an excess of peptide (4). Consequently, chemical shifts in the presence and absence of bicelles are highly similar, and NMR assignments could be readily transferred between the two types of samples. Lipid resonances were assigned from the literature (36) and are summarized in **Fig. S11**. The trNOE correlations for NMR structure calculations were obtained from 2D-NOESY (t_m_ = 150 ms) experiments. The optimal t_m_ value was chosen from the linear portion of a NOE buildup curve (**Fig. S18**), to minimize spin diffusion effects (32, 100).

### NMR structure calculation and analysis

Quantification of NOESY peak intensities and NMR structure calculations were done with the programs CCPN Analysis (101) and Xplor-NIH (102), respectively, on the NMRbox platform (103). The input data for NMR structure calculations were ROE (free peptide) or trNOE distance constraints (bound peptide), together with broad dihedral restraints of φ = −90 ± 70^°^, ψ = 60 ±120° to maintain backbone torsional angles of the two central residues in common regions of Ramachandran space. NMR structures were calculated using simulated annealing and refinement protocols starting from 80 initial conformers with random backbone φ, ψ dihedral angles. For each peptide, the 20 lowest energy structures with no violations outside the thresholds specified in **Tables S2** and **S3** were kept for analysis. Structure #1 in the resulting NMR structure bundles is the closest to the ensemble mean. Electrostatic surfaces for membrane-bound peptides were calculated with the APBS program (104).

Least squares structure superpositions of the NMR bundles were calculated with the FIT routine of the program MolMol (105). To estimate the limiting maximum RMSD for tetrapeptides with unrelated structures we used two approaches. In the first, we randomly extracted 10 tetrapeptide fragments from high-resolution protein structures in the PDB. The backbone RMSD was ∼1Å, while the heavy atom RMSD could not be calculated because of different side chains. In a second approach we calculated SS-peptide structures without any experimental distance or dihedral restraints. We obtained backbone RMSDs of 1 to 2 Å, and heavy atom RMSDs of 3 to 5 Å.

### Cell culture

Human renal epithelial cells (HK-2) and human retinal pigment epithelial cells (ARPE-19) were obtained from ATCC. HK-2 cells were cultured in DMEM (Dulbecco’s Modified Eagle’s Medium) containing 1 g/L glucose and 10% fetal bovine serum (FBS), 100 units/ml penicillin, and 100 μg/ml streptomycin. ARPE-19 cells were grown in DMEM/F12 medium containing 1 g/L glucose and 10% fetal bovine serum (FBS), 100 units/ml penicillin, and 100 μg/ml streptomycin.

### Cell uptake and mitochondrial localization of CL-binding peptides

Cell uptake of SS-31 and SPN10 was determined using N-biotinylated SS-31 and SPN10 and detected by streptavidin binding. After 3 days serum deprivation to deplete endogenous biotin, cells were treated with 1 μM biotinylated peptides for 1h before they were fixed with 4% PFA and incubated with Streptavidin-AlexaFluor 594 antibody (Jackson ImmunoResearch, West Grove, PA) and Hoechst 33342 (Novus Biologicals, Centennial, CO). Images were obtained with a Nikon Eclipse Ti2 fluorescent microscope using a 100X objective.

### Cell viability and cellular ATP after 7 days serum deprivation

Cells were seeded in 96-well culture plates at an initial density of 5 × 10^3^ cells. On the day of the experiment, FBS was removed from the culture medium and cells were incubated in serum-free DMEM alone (control group) or serum-free DMEM containing 10 nM peptide analogs for 7 days. All treatments were carried out with N=4-6 in each experiment. Cell viability was measured by resazurin fluorescence (alamarBlue) from Thermo Fisher (Waltham, MA). ATP was measured using the ApoSENSOR ATP Bioluminescence Assay Kit (BioVision, Milpitas, CA). Luminescence was measured using a microplate reader (SpectraMax iD3, Molecular Devices, San Jose, CA). Results in each experiment were normalized to control and all data are presented as mean ± SEM from 4 experiments. Differences among groups were compared by one-way ANOVA. *Post hoc* analyses were carried out using Tukey’s multiple comparisons test.

### Statistical analyses and scientific rigor

All means reported represent a minimum of n=3 independently prepared sample replicates and are reported as means ± standard deviation (SD) or standard error of the mean (SEM), as appropriate. Statistical analyses were performed using one-way ANOVAs or Wilcoxon rank sum tests as indicated. Differences among sample populations were considered significant at *P* < 0.05.

## Acknowledgements

This work was supported by the National Institutes of Health (Grant R01-AG065879 to N.N.A. and Grant R35-GM119762 to E.R.M.), by The Barth Syndrome Foundation (2020 Idea Grant to N.N.A.) and by a charitable contribution from the Social Profit Network (to N.N.A.)

