## Supplemental Information for "Structure-Activity Relationships in the Design of Mitochondria-Targeted Peptide Therapeutics"

### Table of Contents

**Supplementary Figures** **Page**


Figure S1. NMR cross-relaxation experiments for the SS-31 peptide 1
Figure S2. RMSD of peptides in solution from MD trajectories 2
Figure S3. Representative sections of 2D NOESY spectra 3
Figure S4. Electrostatic surfaces of peptides in the bound state 4
Figure S5. SS-31 side chain insertion depths from MD simulations 5
Figure S6. SS-20 side chain insertion depths from MD simulations 6
Figure S7. SPN4 side chain insertion depths from MD simulations 7
Figure S8. SPN10 side chain insertion depths from MD simulations 8
Figure S9. Comparison of peptide side chain insertion depths from MD simulations 9
Figure S10. NOE restraint violations from peptide-bilayer MD simulations 10
Figure S11. Assignments of bicelle lipid NMR signals 11
Figure S12. trNOEs between peptides and lipids ` 12
Figure S13. Time courses of SASA measurements from MD trajectories 13
Figure S14. Bilayer thickness and area per lipid measurements from MD simulations 14
Figure S15. Lipid radial distribution profiles from MD simulations 15
Figure S16. Peptide analog binding footprints and membrane area coverage 16
Figure S17. Blocked standard error for several analyses from MD simulations 17
Figure S18. trNOE builidup curves 18


**Supplementary Tables** **Page**


Table S1. NMR chemical shift assignments for peptides evaluated 19
Table S2. Statistics for the 20 lowest-energy NMR structures of peptides in their free states 21
Table S3. Statistics for the 20 lowest-energy NMR structures of peptides in their bicelle-bound states 22
Table S4. RMSD and $R_g$ values for NMR measurements and MD simulations 23
Table S5. Improved structural agreement between MD and NMR when imposing NOE Restraints 24

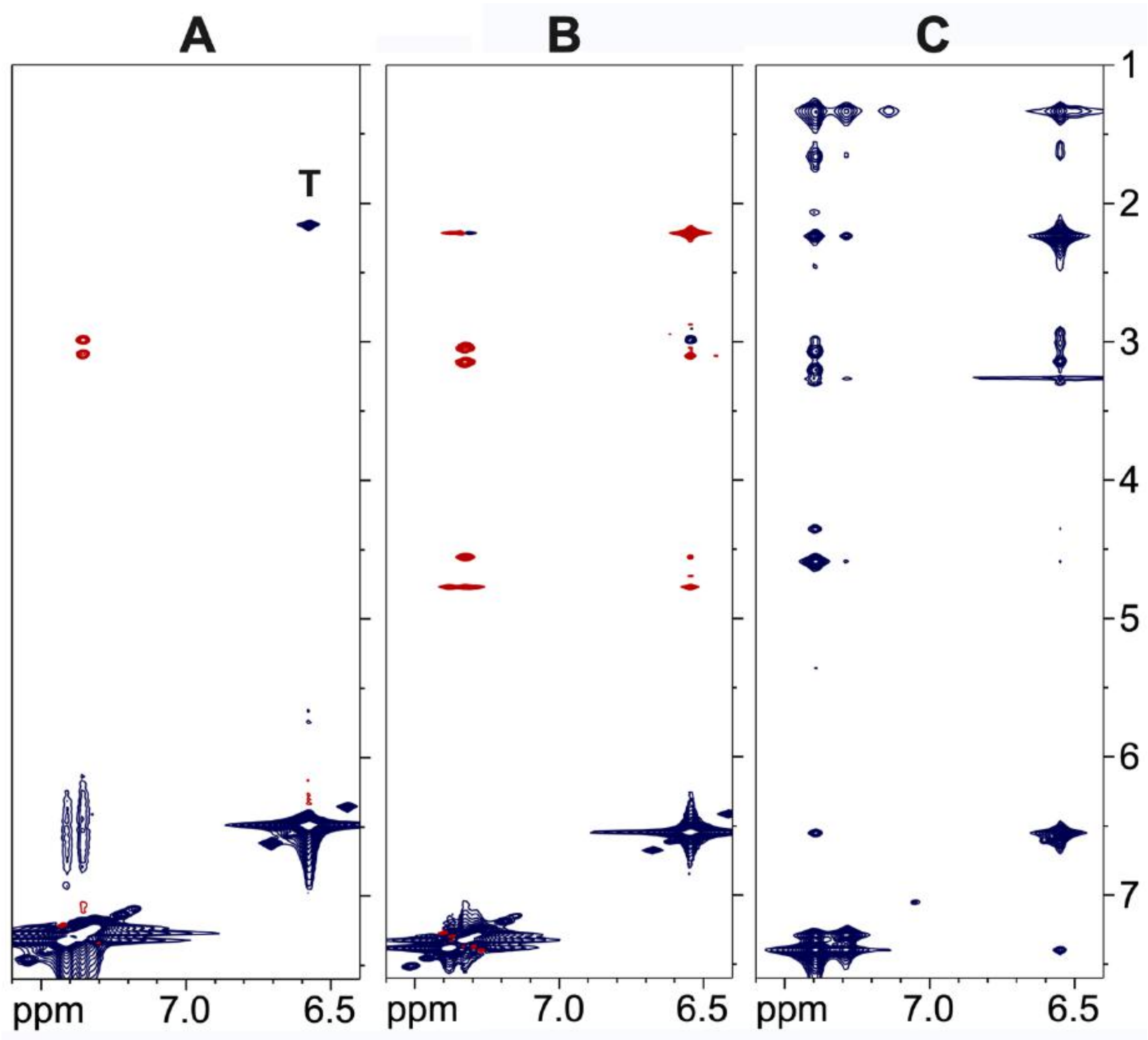


**Figure S1. NMR cross-relaxation experiments for the SS-31 peptide.** (**A**) NOESY of the free peptide at a 10 mM concentration. There are only a small number of positive NOEs (red) due to the small size of the peptide. The peak marked "T" is a TOCSY artifact due to coupling of the δ methyl protons and ε aromatic protons of the 2,6-dimethyl tyrosine residue at position 2. (**B**) ROESY of the free peptide at a 10 mM concentration showing positive crosspeaks. (**C**) trNOESY of the bicelle-bound peptide at a 0.75 mM peptide concentration, showing a large number of strong negative trNOEs (blue) that are transferred from the large MW peptide-bicelle complex to the free peptide.

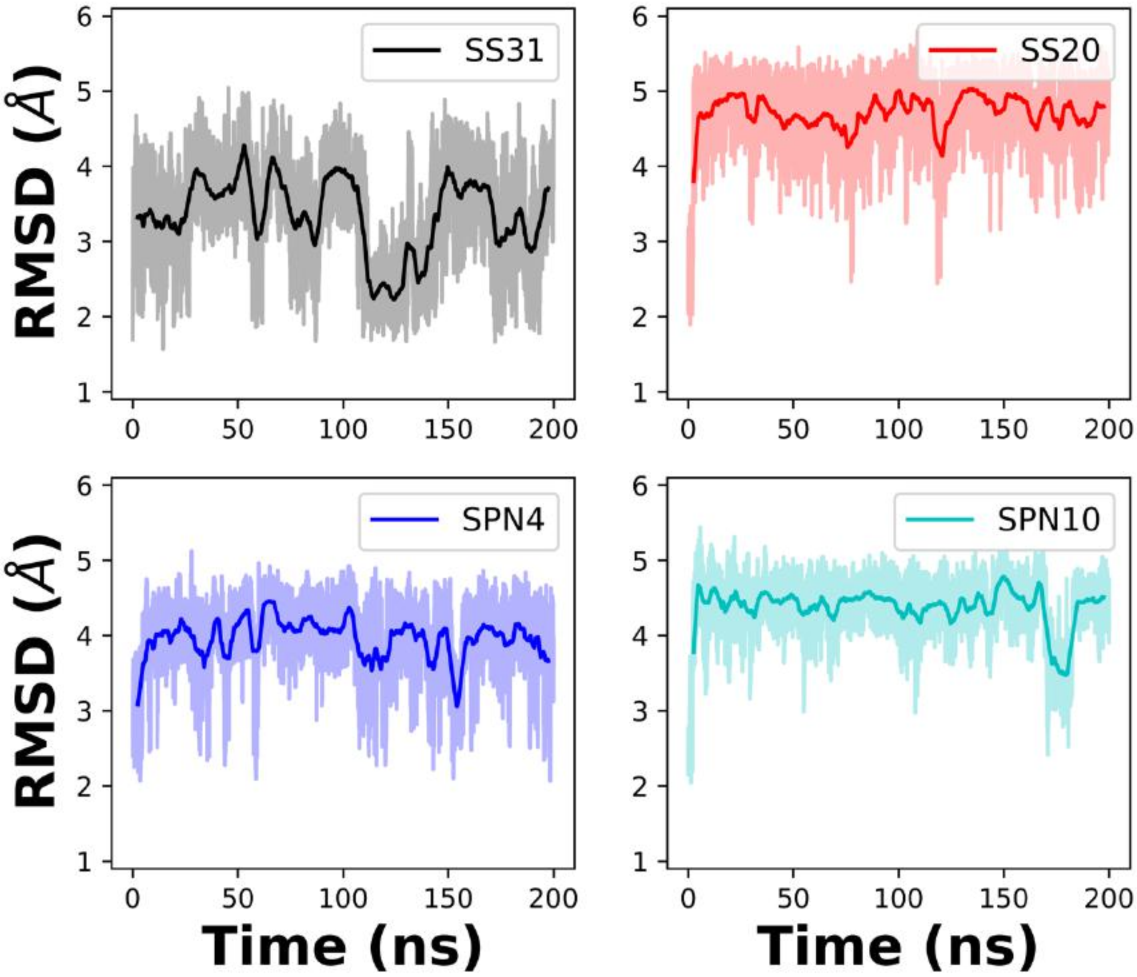


**Figure S2. RMSD of peptides in solution from MD trajectories.** Comparison of the RMSD of each peptide analog in solution over the 200 ns simulation time. The NMR structures in solution were used as initial structures.

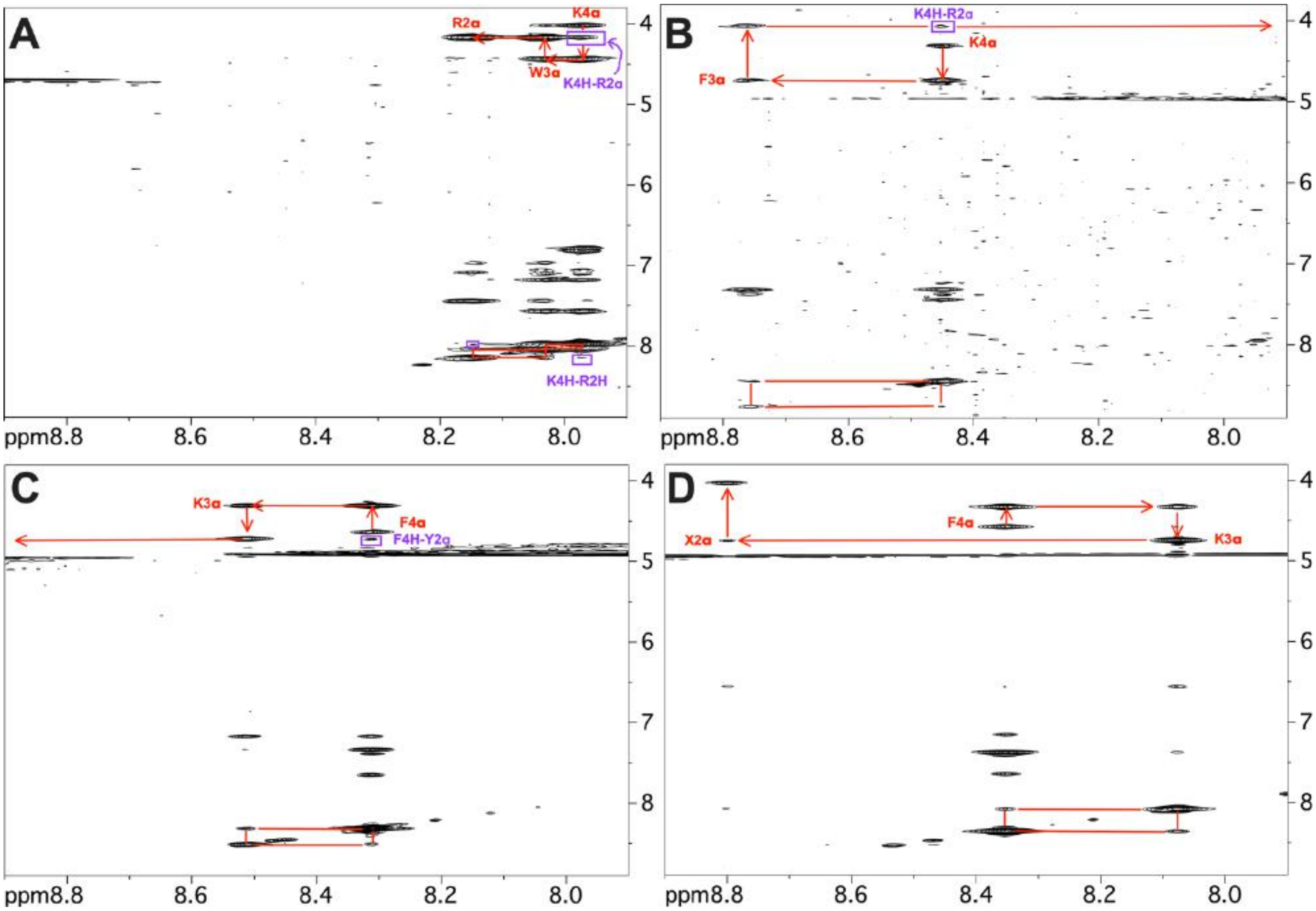


**Figure S3. Representative sections of 2D NOESY spectra.** Portions of 2D NOESY spectra collected for the four peptide analogs showing sequential assignments (red), and NOEs indicative of reverse turn conformations (purple). (**A**) SPN10, (**B**) SS-20, (**C**) SPN10, (**D**) SS-31. Three of the peptides show dαN(i,i+2) NOEs that are consistent with reverse turn but not extended structures (purple). The SPN10 peptide has an additional weak dNN(i,i+2) NOE consistent with a turn. In addition to these backbone connectivities, many additional side chain NOEs also support the turn structures. In contrast, the SS-31 peptide only shows sequential backbone NOEs and has an extended structure. The NOEs to resonances between 6 and 7 ppm are to aromatic side chains in the peptides. As expected, the amino protons of the first residue in each peptide are not seen due to fast solvent exchange. Fast solvent exchange also causes the amide proton of the second residue to be extremely weak or not seen in some of the spectra. In the SS-31 peptide "X2" indicates the 2,6 dimethyl tyrosine residue at position 2 in the sequence.

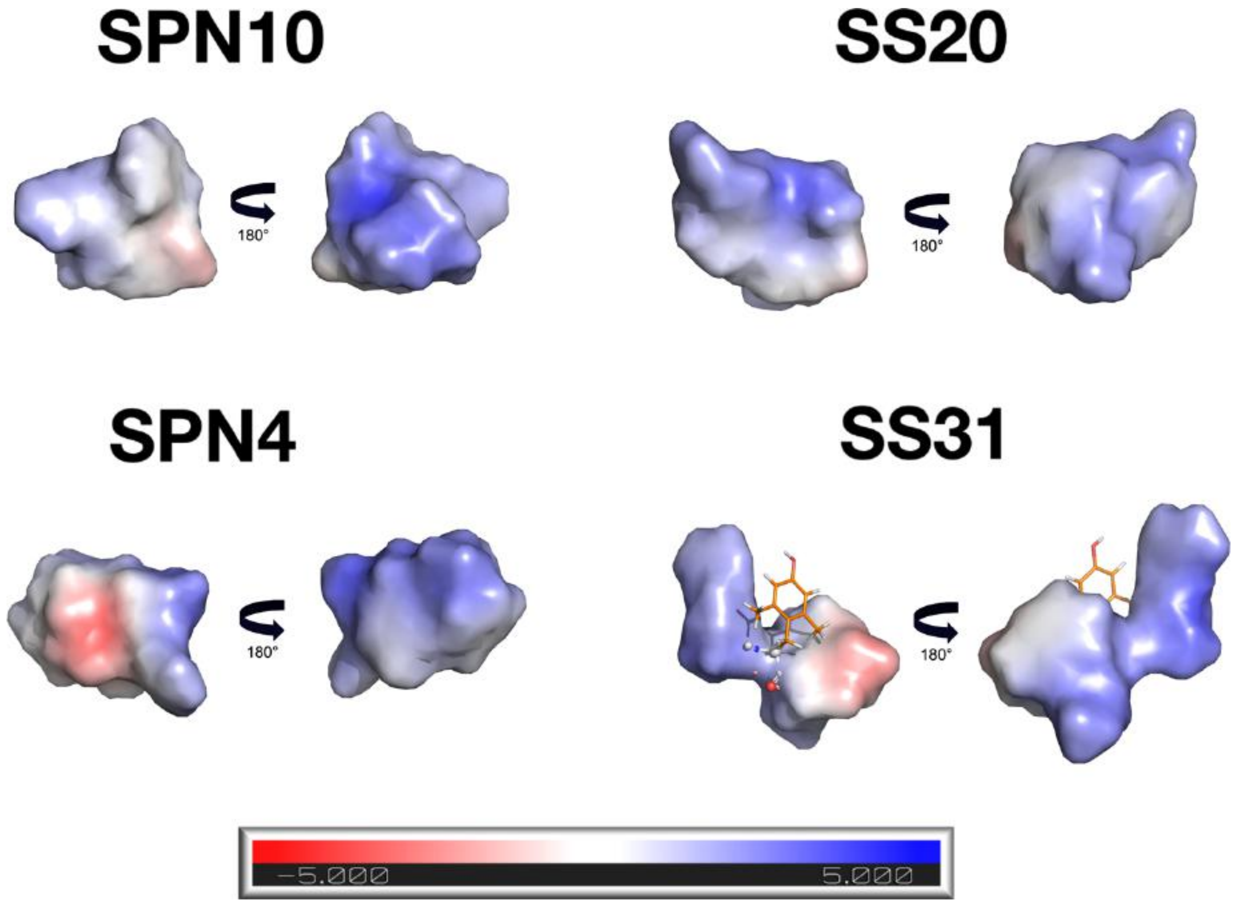


**Figure S4. Electrostatic surfaces of peptides in the bound state.** Electrostatic surfaces for the peptides in the bound state were calculated with the APBS program (**ref. 104**). For each peptide the first view corresponds to that in **Fig. 3** and shows a mostly neutral surface. The second is after a 180° rotation along y, and shows a predominately positively-charged side of the molecule. For the SS-31 peptide the electrostatic potential was not calculated for dimethyl tyrosine since it is a non-standard amino acid.

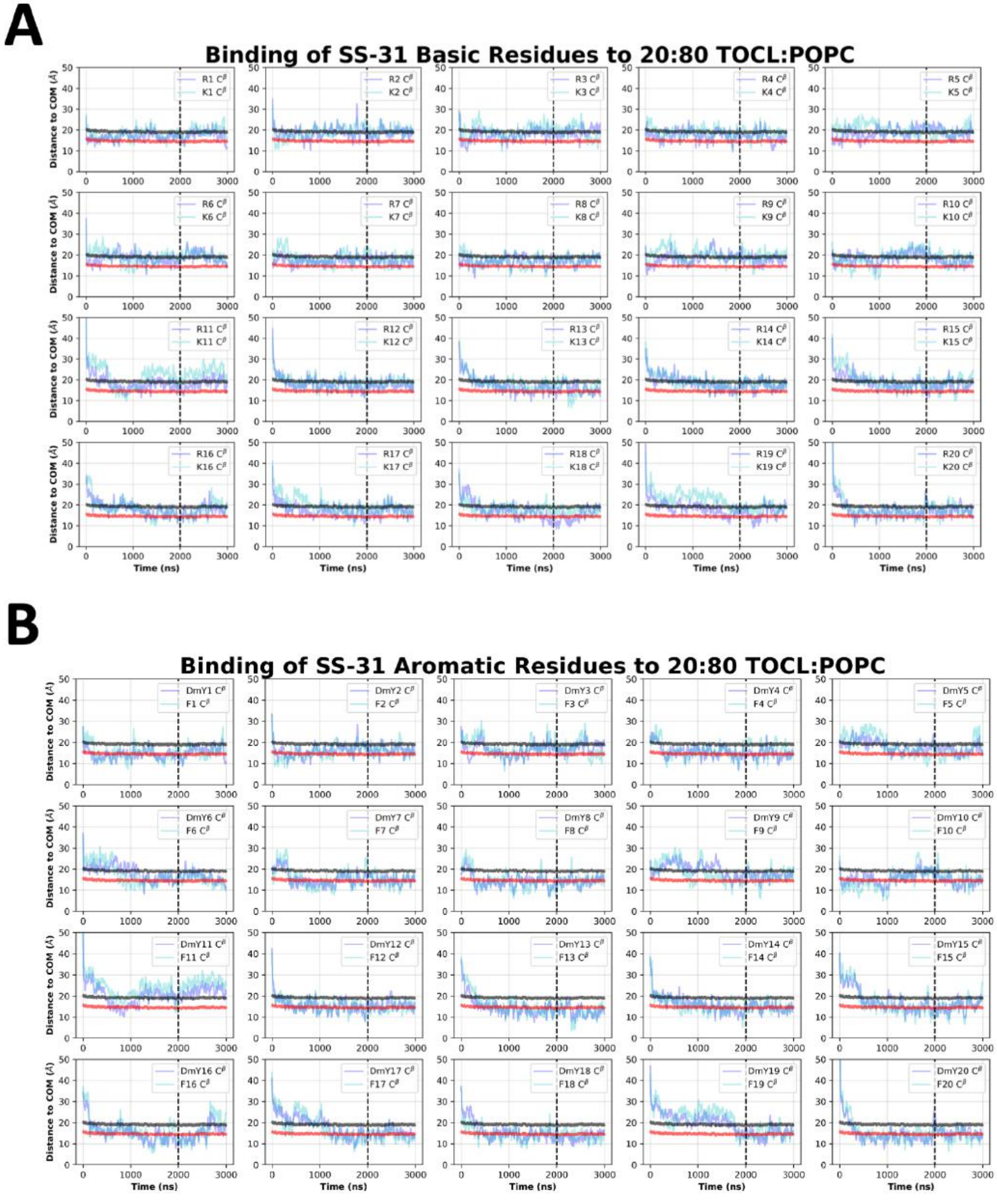


**Figure S5. SS-31 side chain insertion depths from MD simulations.** Time course profiles shown for individual SS-31 basic (**A**) and aromatic (**B**) side chain positions relative to the bilayer. Positions along the z-axis were determined as the distance between the $C^{\beta}$ atom on each side chain and the calculated COM of the membrane. The black and red horizontal lines, respectively, show the average position of all lipid phosphates and lipid ester carbons relative to the membrane COM. Colored lines show the z coordinates ($Z^{pos}$) of individual side chains smoothed over 10 ns intervals. The black vertical dotted line indicates when NMR restraints were implemented.

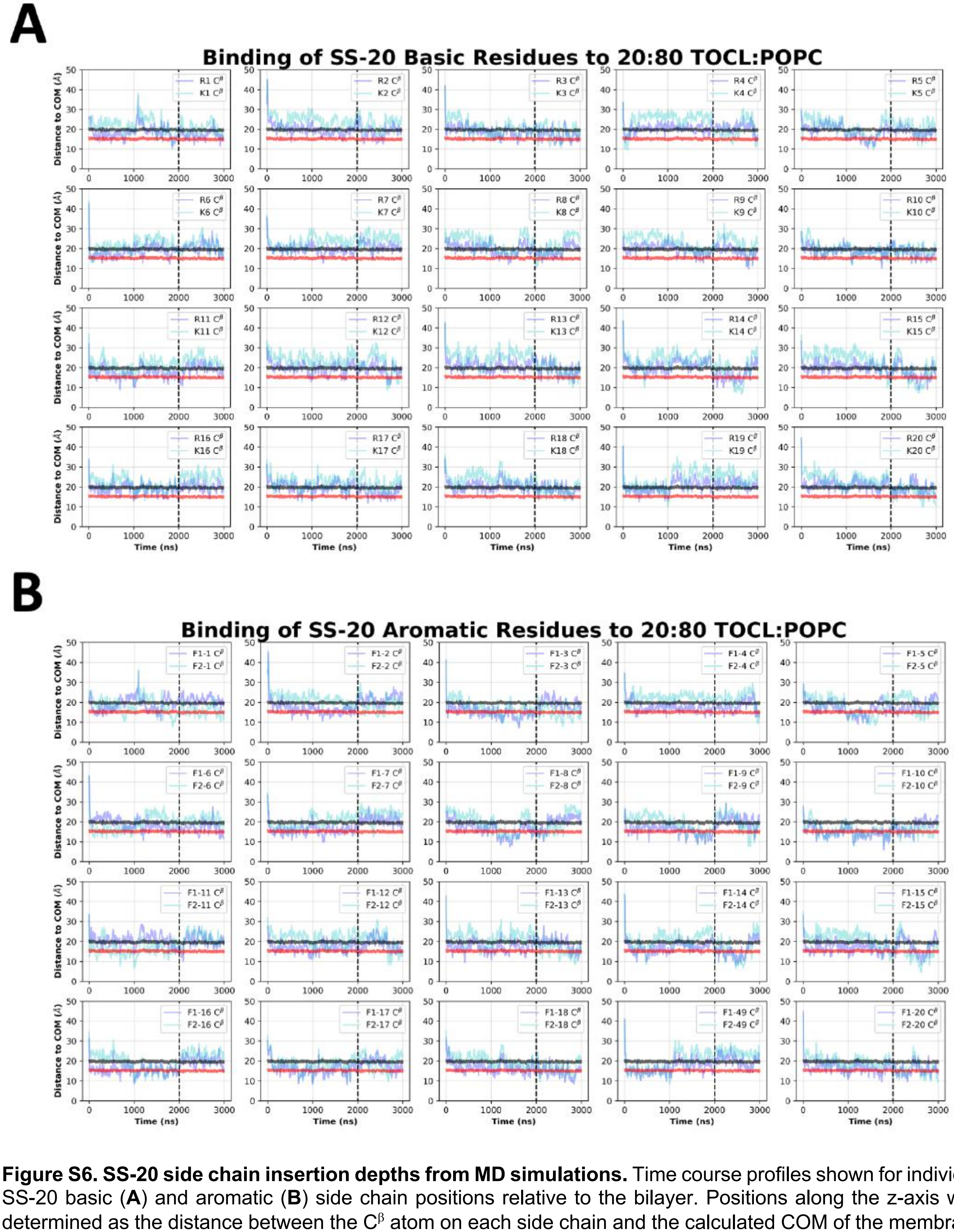


**Figure S6. SS-20 side chain insertion depths from MD simulations.** Time course profiles shown for individual SS-20 basic (**A**) and aromatic (**B**) side chain positions relative to the bilayer. Positions along the z-axis were determined as the distance between the $C^{\beta}$ atom on each side chain and the calculated COM of the membrane. The black and red horizontal lines, respectively, show the average position of all lipid phosphates and lipid ester carbons relative to the membrane COM. Colored lines show the z coordinates ($Z^{pos}$) of individual side chains smoothed over 10 ns intervals. The black vertical dotted line indicates when NMR restraints were implemented.

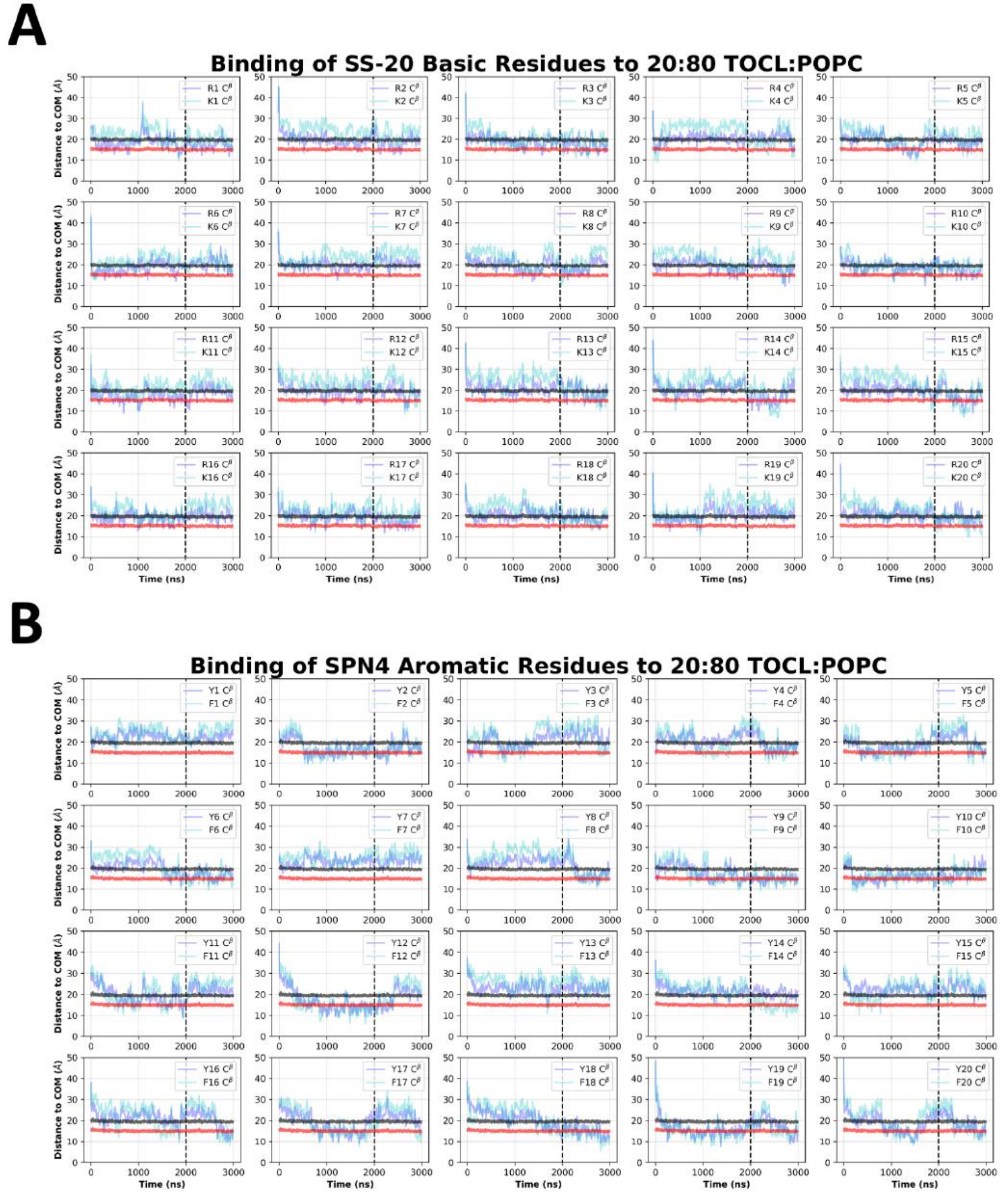


**Figure S7. SPN4 side chain insertion depths from MD simulations.** Time course profiles shown for individual SPN4 basic (**A**) and aromatic (**B**) side chain positions relative to the bilayer. Positions along the z-axis were determined as the distance between the $C^{\beta}$ atom on each side chain and the calculated COM of the membrane. The black and red horizontal lines, respectively, show the average position of all lipid phosphates and lipid ester carbons relative to the membrane COM. Colored lines show the z coordinates ($Z^{pos}$) of individual side chains smoothed over 10 ns intervals. The black vertical dotted line indicates when NMR restraints were implemented.

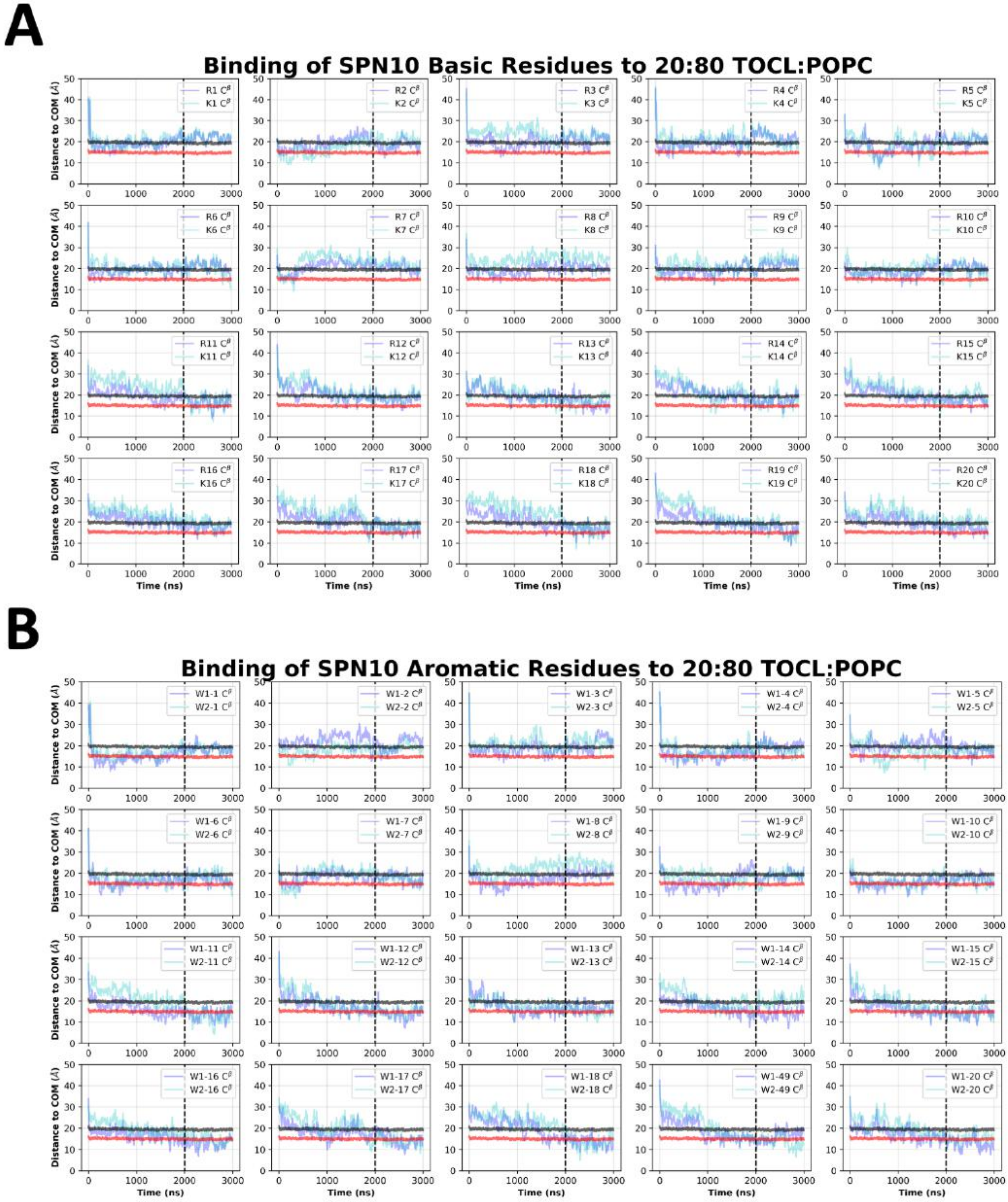


**Figure S8. SPN10 side chain insertion depths from MD simulations.** Time course profiles shown for individual SPN10 basic (**A**) and aromatic (**B**) side chain positions relative to the bilayer. Positions along the z-axis were determined as the distance between the $C^{\beta}$ atom on each side chain and the calculated COM of the membrane. The black and red horizontal lines, respectively, show the average position of all lipid phosphates and lipid ester carbons relative to the membrane COM. Colored lines show the z coordinates ($Z^{pos}$) of individual side chains smoothed over 10 ns intervals. The black vertical dotted line indicates when NMR restraints were implemented.

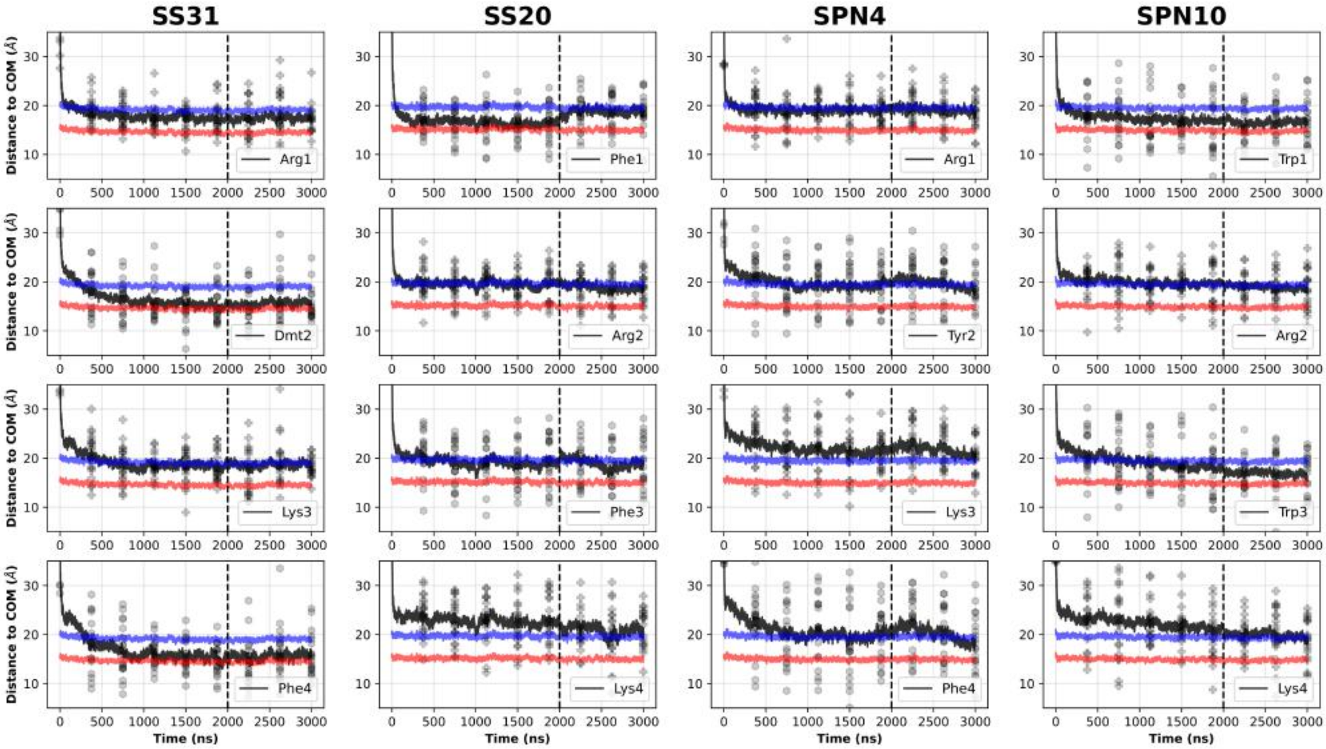


**Figure S9. Comparison of peptide side chain insertion depths from MD simulations.** Time course profiles summarizing the profiles shown in **Fig. S5-S8**, showing the peptide basic (*plus sign*) and aromatic (*hexagon*) side chain positions relative to the bilayer over time. Positions along the z-axis were determined as the distance between the $C^{\beta}$ atom on each side chain and the calculated COM of the membrane. The black and red horizontal lines respectively show the average position of all lipid phosphates and lipid ester carbons relative to the calculated COM of the membrane for each peptide system. Black traces show the average z coordinates ($Z^{pos}$) across all side chains (20 peptides/system) smoothed over 10 ns intervals. The black vertical dotted line indicates when NMR restraints were implemented.

In general, the residues of SS-31 buried the deepest, followed by SPN10, SS-20, and SPN4. Whereas for SS-31 the lysine buried below the lipid phosphates, the lysine of all other peptides were at or above this plane. A similar trend was established amongst the peptides for the arginine moieties, except that all these residues were buried below the plane of the lipid phosphates. Additionally, the same trend applied to the burial depths of the aromatics; perhaps most notably, SPN4 was the only peptide to have aromatics positioned above the plane of the lipid phosphates.

**A**

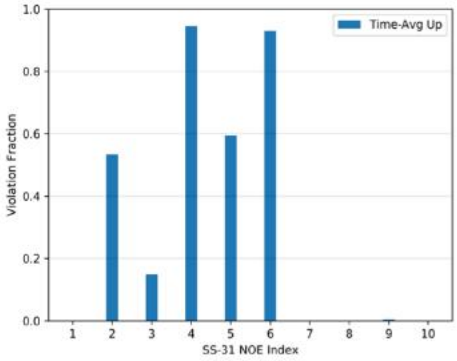


**B**

| Variant: | SS-31 | | | | | | | |
|---|---|---|---|---|---|---|---|---|
| Sequence: | 1\|ARG | 2\|DmTYR | 3\|LYS | 4\|PHE | | | | |
| NOE INDEX | Atom 1 | | Atom 2 | | TYPE | d | $d_{minus}$ | $d_{plus}$ |
| index 1 | Dmt2 HE# | | to Phe4 HE# | | i to i+2 | 4.561 | 0.912 | 0.912 |
| index 2 | Dmt2 HB# | | to Phe4 HN | | i to i+2 | 4.264 | 0.853 | 0.853 |
| index 3 | Dmt2 HE# | | to Phe4 H2/HT2 | | i to i+2 | 4.122 | 0.824 | 0.824 |
| index 4 | Phe4 HN | | to Dmt2 HA | | i to i+2 | 4.219 | 0.844 | 0.844 |
| index 5 | Phe4 HD# | | to Dmt2 HA | | i to i+2 | 3.443 | 0.689 | 1.689 |
| index 6 | Phe4 HN | | to Dmt2 HX# | | i to i+2 | 3.390 | 0.678 | 0.678 |
| index 7 | Phe4 HE# | | to Dmt2 HX# | | i to i+2 | 3.646 | 0.729 | 0.729 |
| index 8 | Phe4 HD# | | to Dmt2 HX# | | i to i+2 | 3.052 | 0.610 | 1.110 |
| index 9 | Phe4 HD# | | to Dmt2 HB1 | | i to i+2 | 3.993 | 0.799 | 1.299 |
| index 10 | Phe4 HA | | to Dmt2 HX# | | i to i+2 | 3.575 | 0.715 | 1.715 |

**C**

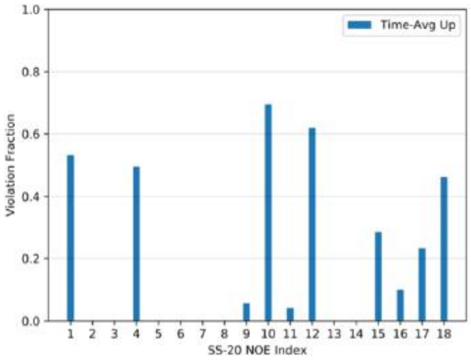


**D**

| Variant: | SS-20 | | | | | | | |
|---|---|---|---|---|---|---|---|---|
| Sequence: | 1\|PHE | 2\|ARG | 3\|PHE | 4\|LYS | | | | |
| NOE INDEX | Atom 1 | | Atom 2 | | TYPE | d | $d_{minus}$ | $d_{plus}$ |
| index 1 | Phe3 HN | | to Phe1 HA | | i to i+2 | 3.581 | 0.716 | 0.716 |
| index 2 | Lys4 HN | | to Arg2 HA | | i to i+2 | 4.663 | 0.933 | 0.933 |
| index 3 | Phe1 HE# | | to Lys4 HG1 | | i to i+3 | 3.616 | 0.723 | 0.723 |
| index 4 | Lys4 HD2 | | to Phe1 HZ | | i to i+3 | 4.407 | 0.881 | 1.381 |
| index 5 | Lys4 HN | | to Phe1 HA | | i to i+3 | 4.053 | 0.811 | 1.811 |
| index 6 | Lys4 HG2 | | to Phe1 HZ | | i to i+3 | 4.438 | 0.888 | 1.188 |
| index 7 | Lys4 HN | | to Phe1 HB2 | | i to i+3 | 4.716 | 0.943 | 1.443 |
| index 8 | Lys4 HB2 | | to Phe1 HD# | | i to i+3 | 4.212 | 0.842 | 0.842 |
| index 9 | Lys4 H1/HT1 | | to Phe1 HD# | | i to i+3 | 3.303 | 0.661 | 0.661 |
| index 10 | Lys4 HN | | to Phe1 HE# | | i to i+3 | 3.930 | 0.786 | 0.786 |
| index 11 | Lys4 HN | | to Phe1 HD# | | i to i+3 | 3.313 | 0.663 | 0.663 |
| index 12 | Lys4 HD1 | | to Phe1 HZ | | i to i+3 | 4.385 | 0.877 | 0.877 |
| index 13 | Lys4 HD2 | | to Phe1 HZ | | i to i+3 | 4.631 | 0.926 | 0.926 |
| index 14 | Lys4 HD1 | | to Phe1 HD# | | i to i+3 | 4.281 | 0.856 | 0.856 |
| index 15 | Lys4 HD# | | to Phe1 HE# | | i to i+3 | 4.281 | 0.856 | 0.856 |
| index 16 | Lys4 HE# | | to Phe1 HE# | | i to i+3 | 4.289 | 0.857 | 0.857 |
| index 17 | Lys4 HE# | | to Phe1 HZ | | i to i+3 | 3.960 | 0.792 | 0.792 |

**E**

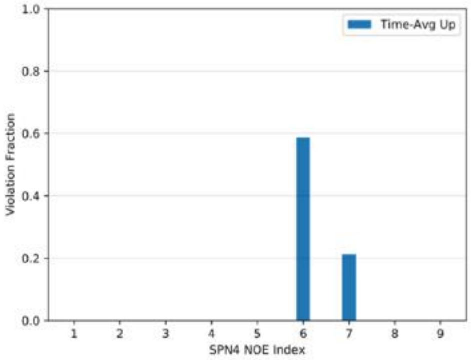


**F**

| Variant: | SPN4 | | | | | | | |
|---|---|---|---|---|---|---|---|---|
| Sequence: | 1\|ARG | 2\|TYR | 3\|LYS | 4\|PHE | | | | |
| NOE INDEX | Atom 1 | | Atom 2 | | TYPE | d | $d_{minus}$ | $d_{plus}$ |
| index 1 | Tyr2 HB2 | | to Phe4 HZ | | i to i+2 | 4.245 | 0.849 | 1.349 |
| index 2 | Tyr2 HB2 | | to Phe4 HD2 | | i to i+2 | 4.495 | 0.899 | 0.899 |
| index 3 | Tyr2 HD1 | | to Phe4 HZ | | i to i+2 | 3.657 | 0.731 | 0.731 |
| index 4 | Phe4 HN | | to Tyr2 HB# | | i to i+2 | 4.335 | 0.867 | 0.867 |
| index 5 | Phe4 HD# | | to Tyr2 HE# | | i to i+2 | 3.817 | 0.763 | 0.763 |
| index 6 | Phe4 HD# | | to Tyr2 HD# | | i to i+2 | 4.460 | 0.892 | 0.892 |
| index 7 | Phe4 HE2 | | to Tyr2 HB2 | | i to i+2 | 4.429 | 0.886 | 0.886 |
| index 8 | Phe4 H1/HT1 | | to Tyr2 HB# | | i to i+2 | 4.179 | 0.836 | 0.836 |

**G**

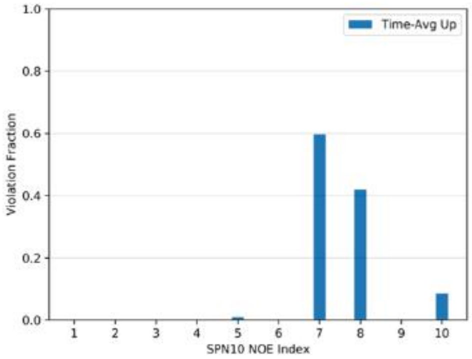


**H**

| Variant: | SPN10 | | | | | | | |
|---|---|---|---|---|---|---|---|---|
| Sequence: | 1\|TRP | 2\|ARG | 3\|TRP | 4\|LYS | | | | |
| NOE INDEX | Atom 1 | | Atom 2 | | TYPE | d | $d_{minus}$ | $d_{plus}$ |
| index 1 | Trp3 HE3 | | to Trp1 HE3 | | i to i+2 | 3.780 | 0.756 | 1.256 |
| index 2 | Lys4 HN | | to Arg2 HN | | i to i+2 | 4.707 | 0.941 | 0.941 |
| index 3 | Lys4 H1/HT1 | | to Arg2 HD# | | i to i+2 | 4.151 | 0.830 | 1.830 |
| index 4 | Trp1 HA | | to Lys4 HN | | i to i+3 | 3.625 | 0.725 | 0.925 |
| index 5 | Arg2 HN | | to Lys4 H1/HT1 | | i to i+3 | 4.071 | 0.814 | 0.814 |
| index 6 | Lys4 HN | | to Trp1 HB# | | i to i+3 | 3.453 | 0.691 | 1.391 |
| index 7 | Lys4 HG# | | to Trp1 HE3 | | i to i+3 | 3.327 | 1.527 | 1.665 |
| index 8 | Lys4 H1/HT1 | | to Trp1 HE3 | | i to i+3 | 2.598 | 0.520 | 1.020 |
| index 9 | Lys4 H1/HT1 | | to Trp1 HB# | | i to i+3 | 4.054 | 0.811 | 1.811 |

**Figure S10. NOE restraint violations form peptide-bilayer MD simulations.** Comparison of the fraction of simulation time in which time-averaged ensembles of peptides violated the upper bound for each NOE restraint for (**A**) SS-31, (**C**) SS-20, (**E**) SPN4, and (**G**) SPN10. Table of NOE restraints imposed in peptide-bilayer MD simulations for (**B**) SS-31, (**D**) SS-20, (**F**) SPN4, and (**H**) SPN10.

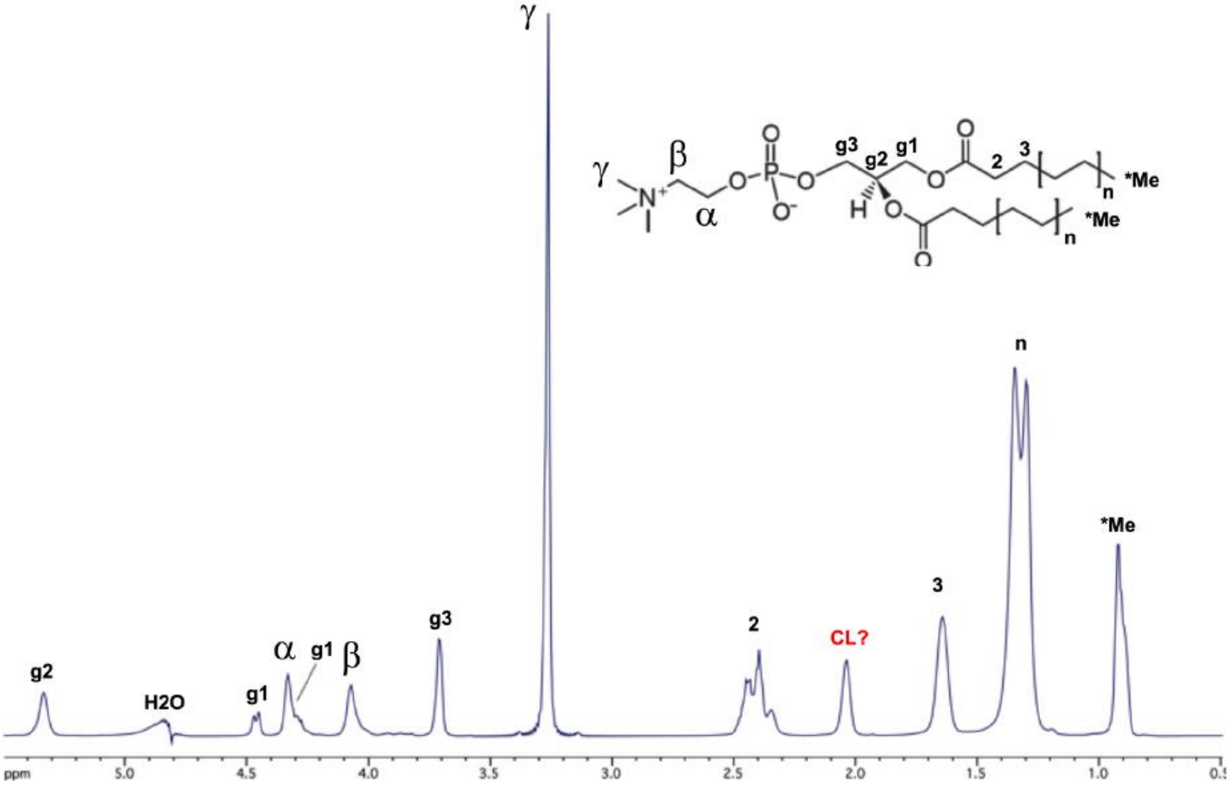


**Figure S11. Assignments of bicelle lipid NMR signals.** The NMR spectrum is of the 150 µM TOCL : 1500 µM POPC : 4500 µM DHPC bicelles (**ref. 4**) in the absence of peptides. The inset shows the chemical structure of POPC, with NMR assignments from a published report (**ref. 36**) indicated on the spectrum. The signal at 2.05 ppm is likely from the CL lipid (possibly the proton on the glycerol *sn*-2 carbon) since it is absent in bicelles that have only the POPC and DHPC lipids. The assignments above were used to interpret NOEs between the peptides and bicelle lipids (see **Fig. S12**).

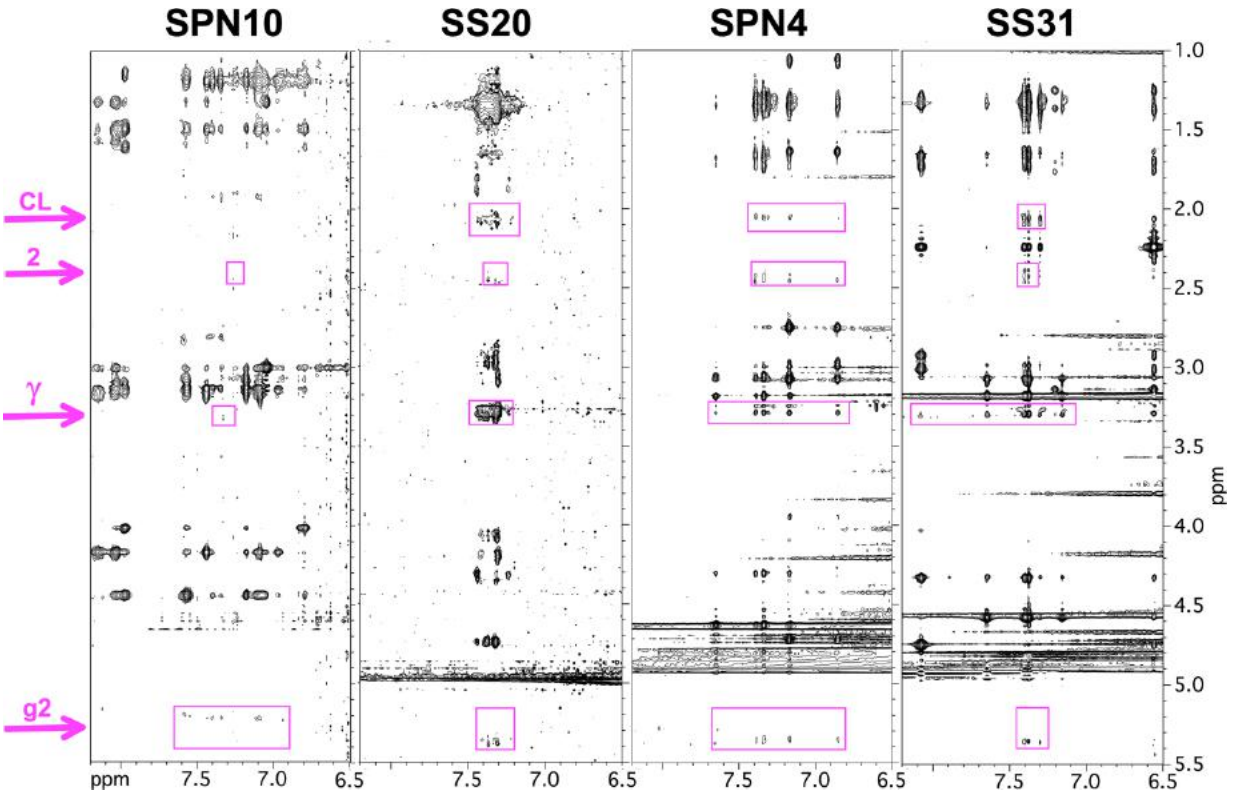


**Figure S12. trNOEs between peptides and lipids.** Sections of the 2D NOESY spectra recorded on peptides in the presence of bicelles, showing NOEs to bicelle lipid protons (highlighted in purple). For NMR assignments see **Fig. S11**. The "CL" signal is specific to cardiolipin and is tentatively assigned to the proton on the 2-glycerol carbon. Only those lipid resonances with no nearby signals from the peptide were selected for analysis. Note that the peptide NOEs for SPN10 are fewer and weaker than for the other peptides.

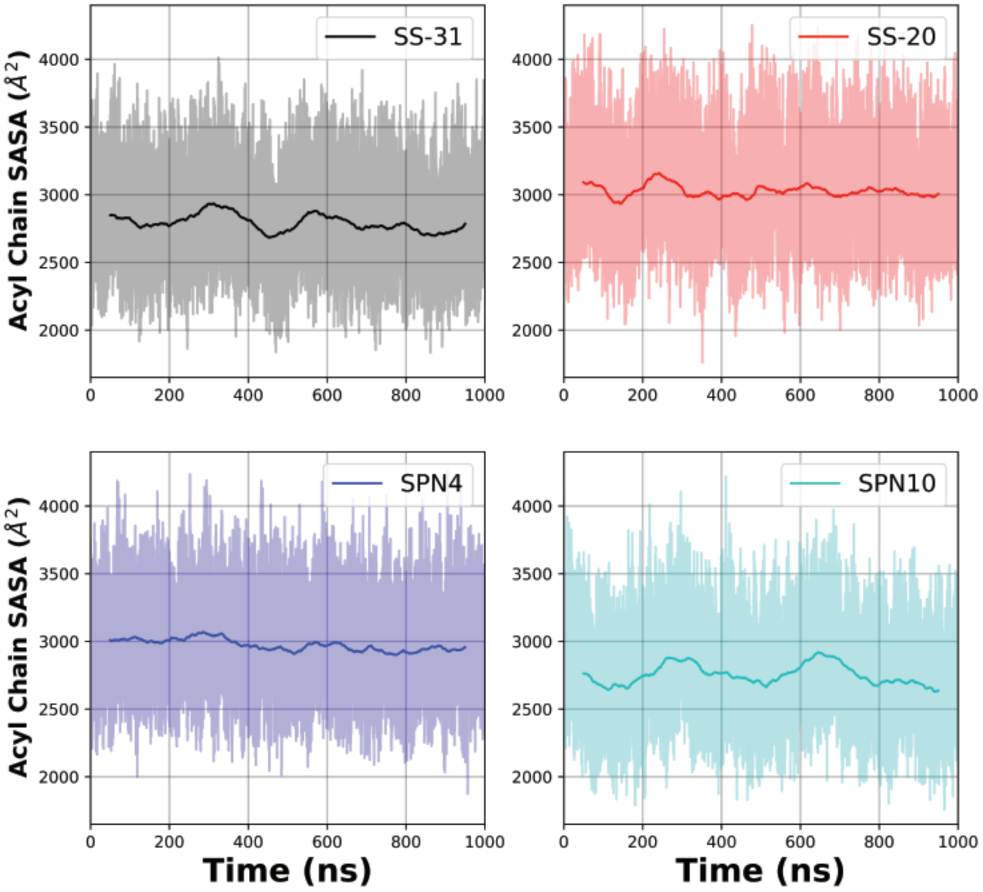


**Figure S13. Time courses of SASA measurements from MD trajectories.** Temporal evolution of the raw (*lighter colored traces*) and smoothed (*thicker traces*, 100 ns rolling average) SASA of bilayer lipid acyl chain region calculated over time for each peptide analog.

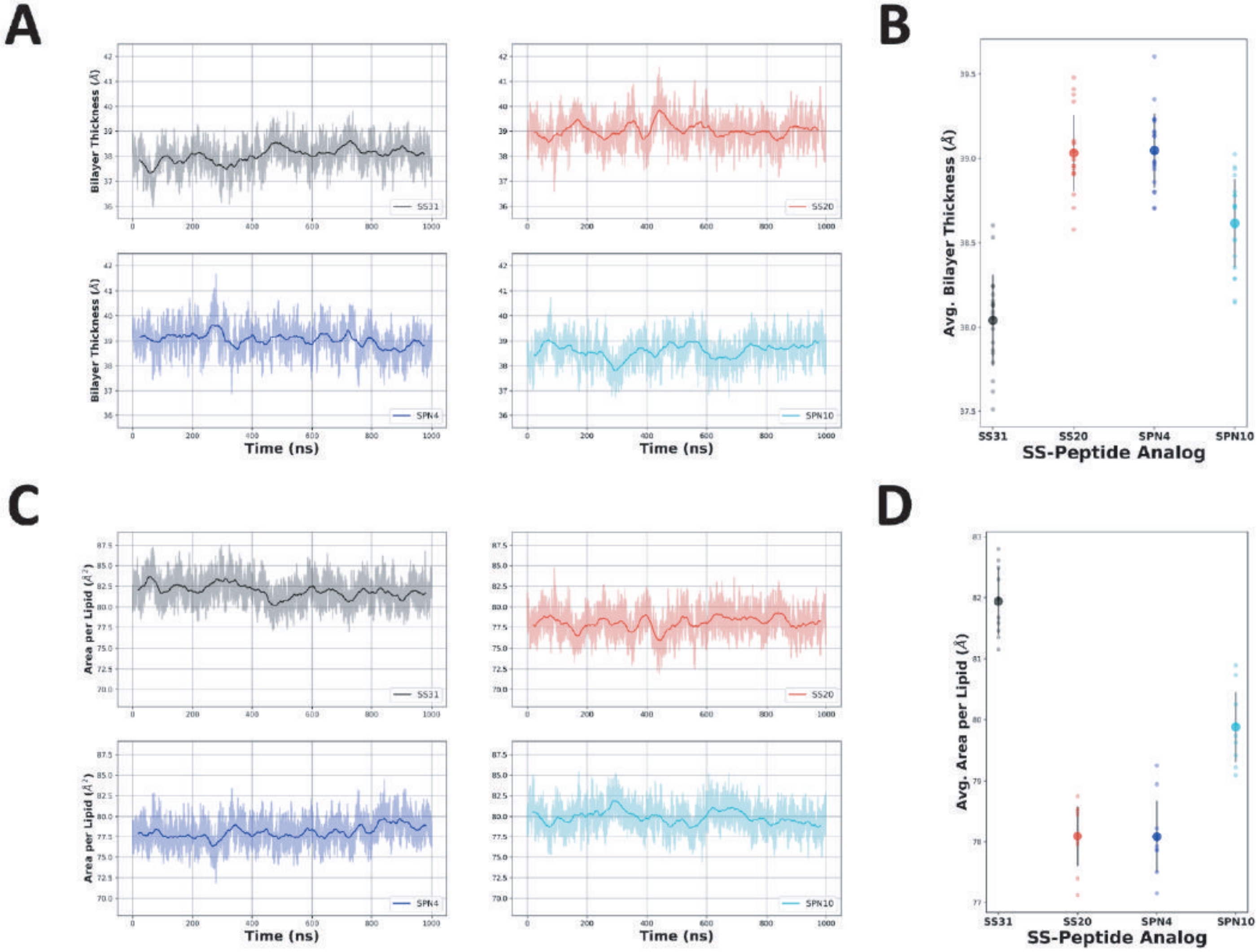


**Figure S14. Bilayer thickness and area per lipid measurements from MD simulations.** (**A**) The raw (*lighter color*) and smoothed (*darker color*, 50 ns rolling average) thickness of each bilayer was calculated as the difference between the average phosphate positions in the upper and lower leaflets over time for each peptide analog. (**C**) The raw (*lighter color*) and smoothed (*darker color*, 80 ns rolling average) area per lipid of each bilayer was calculated over time for each peptide analog. The average bilayer thickness (**B**) and area per lipid (**D**) of each bilayer in the presence of different peptide analogs are shown with standard deviations calculated from 50 ns blocks in the last microsecond of each trajectory.

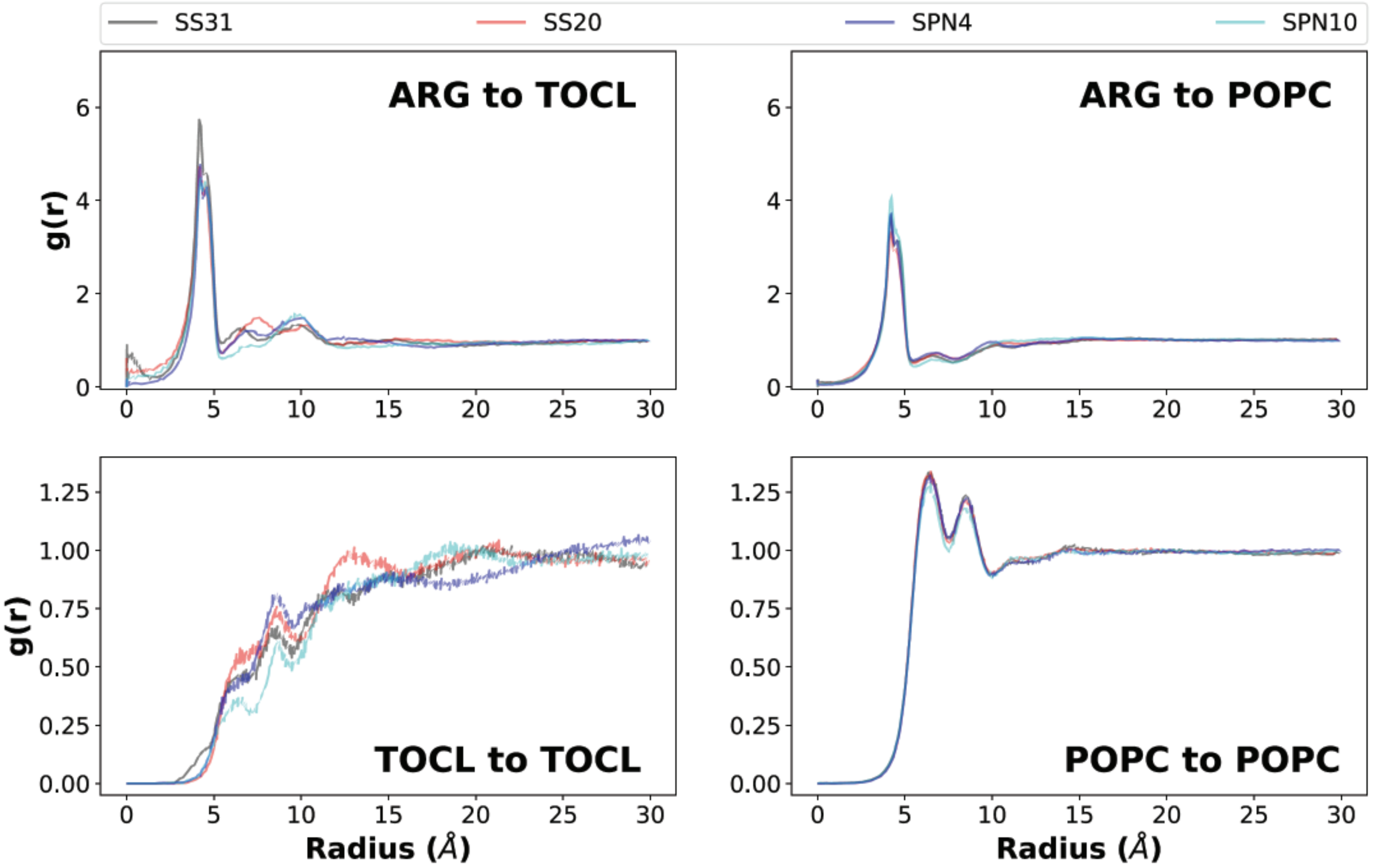


**Figure S15. Lipid radial distribution profiles form MD simulations.** *Upper panels,* Lipid distributions around different peptide analogs. Lateral (x-y plane) radial distributions averaged across both leaflets from the Arg (reference is the Arg-$C^{\zeta}$ atom) to lipid headgroup phosphate atoms (*top left*: TOCL, P1; *top right*: POPC, P). Three distinct radial shells of TOCL and POPC phosphate groups are observed around the Arg-$C^{\zeta}$ atom: a primary shell (radius ~ 4 Å), a secondary shell (radius ~6-8 Å) and a tertiary shell (radius ~9-12 Å). Notably, SS-31 had an additional density in the 0-2 Å range, which agreed with the insertion depth data showing an Arg residue residing below the phosphates on average and can thus overlap in the x-y plane for this radial distribution. *Lower panels*, Lipid-to-lipid distributions in the presence of different peptide analogs. Lateral (x-y) radial distribution profiles of phosphate to phosphate in each lipid group (*bottom left*: TOCL, P1; *bottom right*: POPC, P). Approximately four radial shells of CL phosphate groups are observed around the $P_1$ atom: a primary density (radius ~ 6 Å), as well as secondary (radius ~8-10 Å), tertiary (radius ~12-15 Å), and quaternary (radius ~16-22 Å) densities. For PC phosphate groups, the four radial shells include a primary density (radius 6-7 Å), as well as secondary (radius ~8-9 Å), tertiary (radius ~11-12 Å), and quaternary (radius ~14-15 Å) densities.

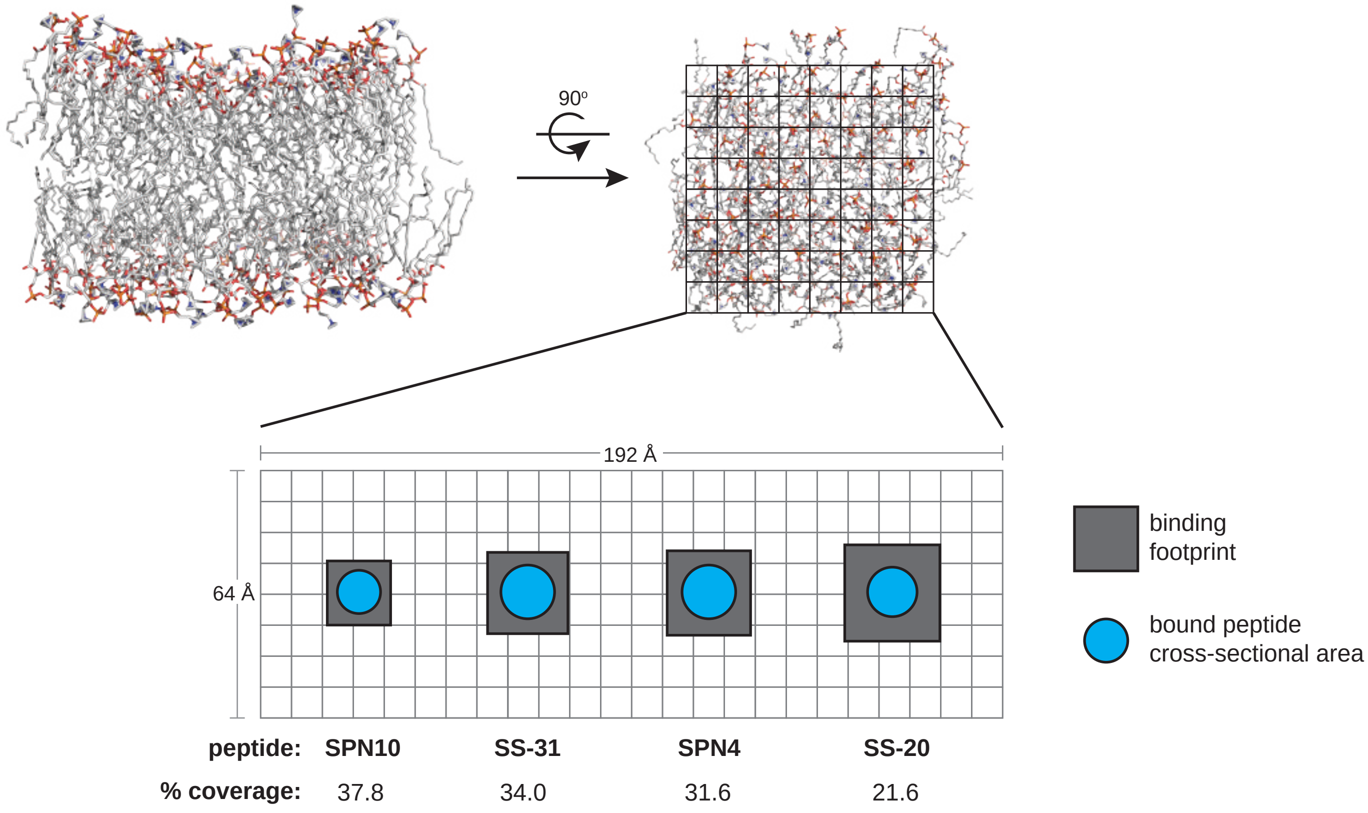


**Figure S16. Peptide analog binding footprints and membrane area coverage.** Schematic depicting the total binding footprint (total membrane area occupied per peptide) and approximate physical cross-sectional area of bound peptide. Top-down view of model membrane is parsed in approximate 8Å x 8Å squares. The peptide-specific binding footprints (*gray boxes*) are estimated from *n* (the lipid:peptide stoichiometry from ITC measurements), equal to 270 Å$^2$ (SPN10), 434 Å$^2$ (SS-31), 466 Å$^2$ (SPN4), and 605 Å$^2$ (SS-20). The peptide-specific cross-sectional areas (*cyan circles*) are estimated from the longest inter-heavy atom distance from our bound NMR structures [11.4 Å (SPN10), 13.7Å (SS-31), 13.7Å (SPN4), and 12.9 Å (SS-20)], which were then used as diameters to estimate a maximal circular cross-sectional area [102 Å$^2$ (SPN10), 147.4 Å$^2$ (SS-31), 147.4 Å$^2$ (SPN4), and 130.7 Å$^2$ (SS-20)]. The % coverage, calculated as cross-sectional area / binding footprint, provides a metric for the amount of membrane area occupied by peptide.

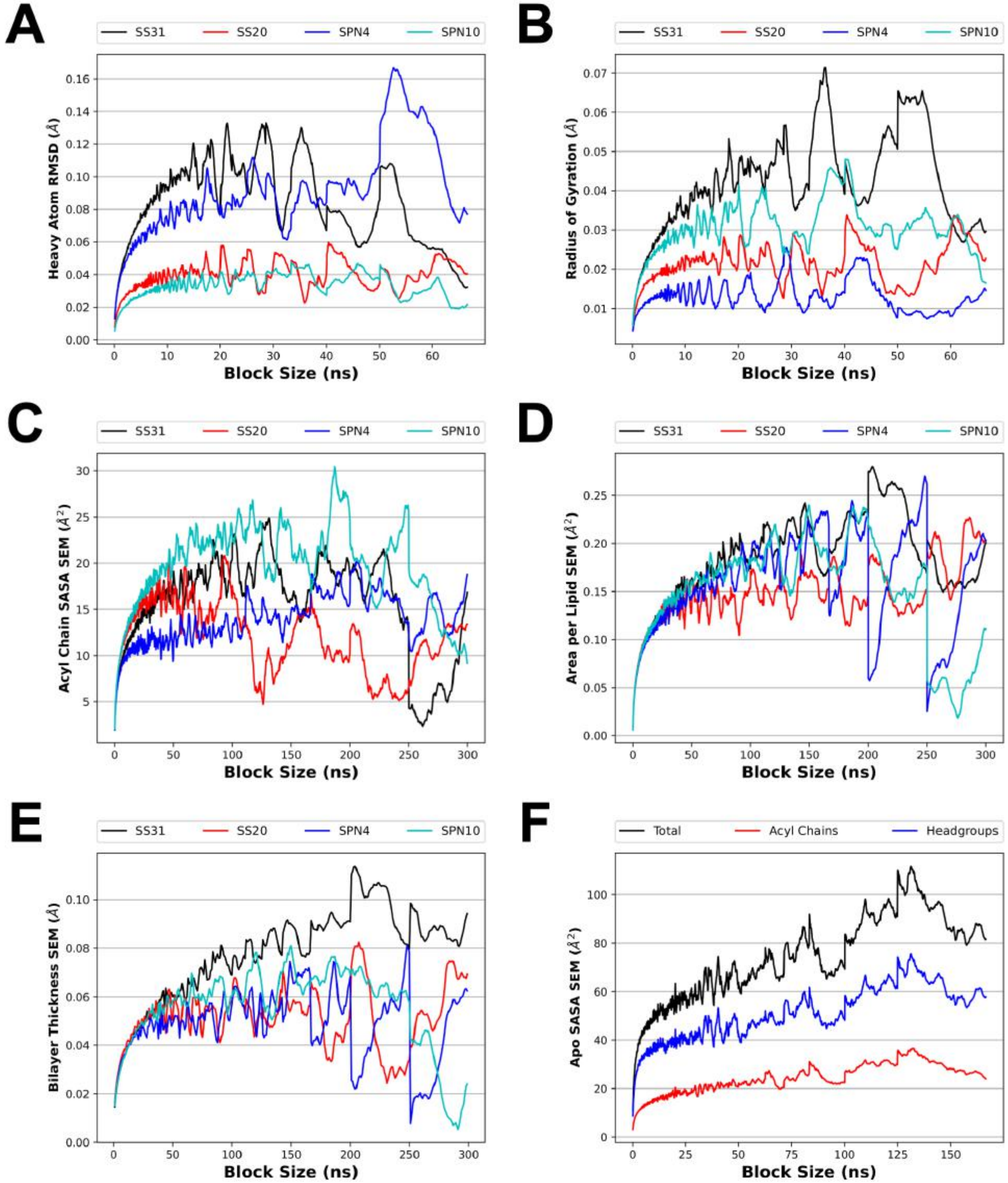


**Figure S17. Blocked standard error for several analyses from MD simulations.** To make optimal use of our sampling from the MD simulations, time course equilibrium data was divided into independent samples. Block averaging was used to select a time interval for which the data would be decorrelated with itself. The blocked standard error was measured for the heavy atom RMSD (**A**) and radius of gyration (**B**) of the MD simulation of a single peptide in solution (**A**, **B**; see **Fig. 1C**). The blocked standard error was also measured for several membrane properties including SASA (**C**,**F**; see **Fig. 4D**), area per lipid (**D**; see **Fig. S14 C,D**), and bilayer thickness (**E**; see **Fig. S14 A,B**). The block sizes selected were 15 ns (**A**,**B**), 25 ns (**F**), and 50 ns (**C**,**D**,**E**). The first block of simulation time was excluded from the block averaging analysis of SASA, APL, and bilayer thickness to account for system relaxation.

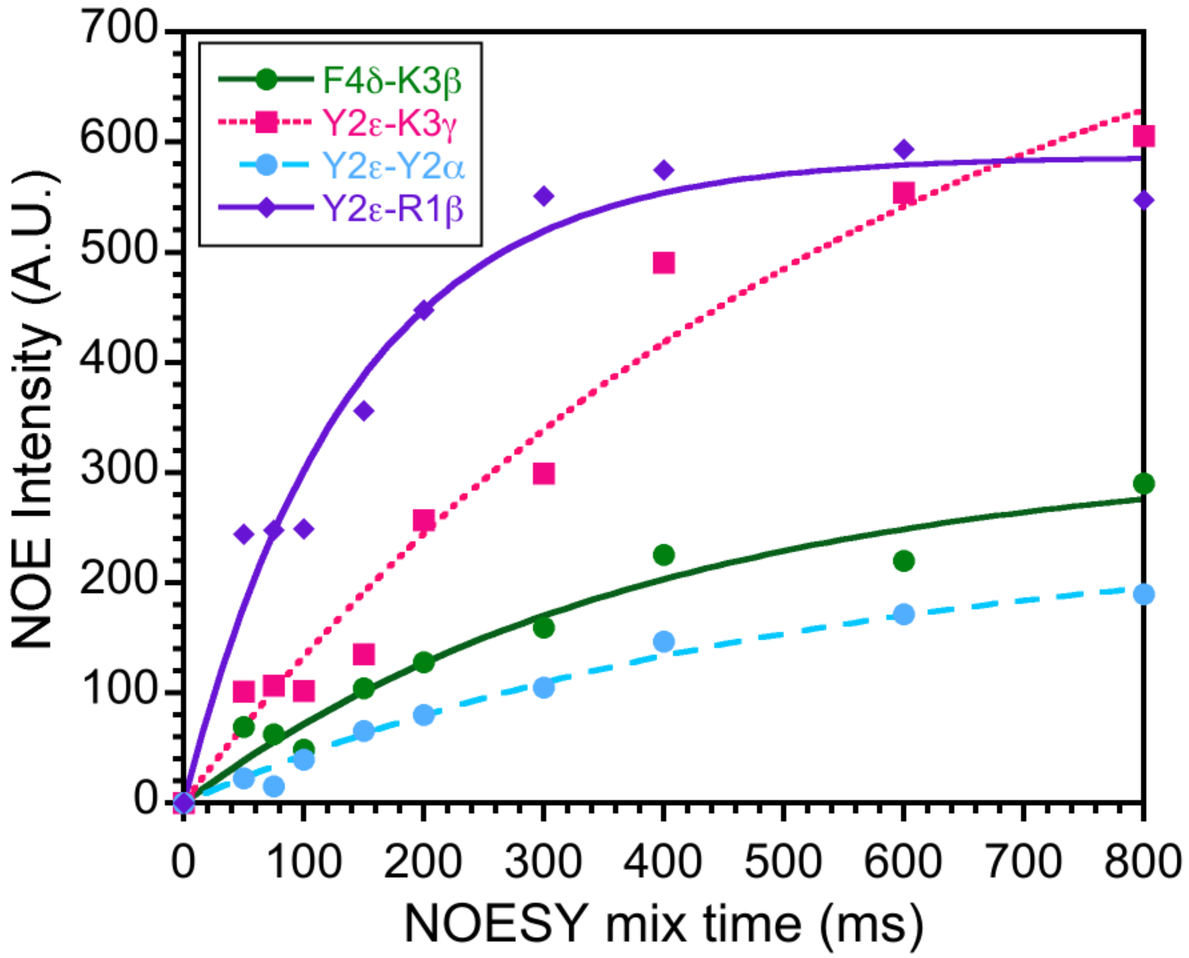


**Figure S18. trNOE buildup curves.** NOESY buildup curves for the bicelle-bound SPN4 sample at 600 MHz. Nine 2D NOESY spectra were collected to monitor the dependence of NOEs on the mixing time. Representative data are shown for the four distance contacts listed in the inset. Based on the data we selected mixing times of 150 ms in the linear portions of the curves, to minimize spin diffusion effects.

**Table S1. NMR chemical shift assignments for the peptides evaluated.**[a,b]

***SPN10***

| | H | N | Hα | Cα | Hβ | Cβ | other |
|---|---|---|---|---|---|---|---|
| Trp1 | - | - | 4.16 | 57.02 | 3.13 | 30.24 | Hδ1 7.06; Hε1 9.99;<br>Hε3 7.44; Hζ2 7.39;<br>Hζ3 7.01; Hη2 7.13;<br>Cδ1 128.61; Cε3 121.21<br>Cζ2 115.27; Cζ3 122.73<br>Cη2 125.34; Nε1 130.32 |
| Arg2 | 8.15 | 123.39 | 4.14 | 56.79 | 1.48,1.54 | 32.00 | Hγ 1.32; Hδ 2.99;<br>Hsc 6.90, 7.05;<br>Cγ 27.47; Cδ 43.86,<br>Nε 96.86 |
| Trp3 | 8.01 | 123.13 | 4.43 | 57.62 | 3.12 | 30.16 | Hδ1 7.16; Hε1 10.09;<br>Hε3 7.56; Hζ2 7.33;<br>Hζ3 7.08; Hη2 7.16;<br>Cδ1 127.99; Cε3 121.52<br>Cζ2 115.34; Cζ3 122.75<br>Cη2 125.37; Nε1 129.51 |
| Lys4 | 7.96 | 124.35 | 4.00 | 56.80 | 1.48,1.59 | 33.68 | Hγ 1.14, Hδ 1.48; Hε 2.78;<br>Cγ 25.01; Cδ 29.87;<br>Cε 42.61 |
| C’$NH_2$ | 6.67,6.77 | 107.45 | | | | | |

***SS20***

| | H | N | Hα | Cα | Hβ | Cβ | other |
|---|---|---|---|---|---|---|---|
| Phe1 | - | - | 4.11 | 57.45 | 3.05, 3.24 | 40.36 | Hδ 7.25; Hε 7.33; Hζ 7.29;<br>Cδ 132.00; Cε 130.59;<br>Cζ 130.07 |
| *arg2* | 7.06 | N.D. | 4.15 | 56.16 | 1.16,1.20 | 30.63 | Hγ 0.72, 0.76; Hδ 2.83;<br>Hsc 6.90, 7.13;Cγ 26.61;<br>Cδ 43.11; Nε 96.80 |
| Phe3 | 8.75 | 122.21 | 4.74 | 57.51 | 2.92, 3.28 | 39.80 | Hδ 7.33; Hε 7.36; Hζ 7.42;<br>Cδ 131.90; Cε 131.62;<br>Cζ 131.83 |
| Lys4 | 8.45 | 123.48 | 4.30 | 56.17 | 1.79, 1.87 | 33.08 | Hγ 1.43, 1.48; Hδ1.67,1.71;<br>Hε 2.99; Cγ 24.92;<br>Cδ 29.10; Cε 41.88 |
| C’$NH_2$ | 7.22,7.43 | 108.63 | | | | | |

***SPN4***

| | H | N | Hα | Cα | Hβ | Cβ | other |
|---|---|---|---|---|---|---|---|
| *arg1* | - | - | 3.96 | 54.35 | 1.66 | 29.84 | Hγ 1.11, 1.12; Hδ 3.01; Cγ 24.19; Cδ 41.79 |
| Tyr2 | 8.86 | 118.64 | 4.73 | N.D. | 2.76, 3.08 | 37.87 | Hδ 7.18; Hε 6.87; Cδ 131.97; Cε 117.02 |
| Lys3 | 8.52 | 123.10 | 4.33 | 55.02 | 1.74, 1.67 | 31.94 | Hγ 1.40, 1.34; Hδ 1.66; Hε 3.00; Cγ 23.21; Cδ 27.85; Cε 40.85 |
| Phe4 | 8.30 | 122.15 | 4.65 | 55.75 | 3.08, 3.21 | 38.37 | Hδ 7.36; Hε 7.38; Hζ 7.30; Cδ 130.74; Cε 130.22; Cζ 128.83 |
| C'$NH_2$ | 7.17,7.65 | 109.37 | | | | | |

***SS31***

| | H | N | Hα | Cα | Hβ | Cβ | other |
|---|---|---|---|---|---|---|---|
| *arg1* | - | - | 3.89 | 55.34 | 1.69 | 31.21 | Hγ 1.29; Hδ 3.13; Cγ 25.55; Cδ 43.09 |
| Tyx2 | 8.81 | 122.40 | 4.75 | 61.92 | 2.96, 3.17 | 33.17 | Hδ 2.24; Hε 6.56; Cδ 21.72; Cε 117.28 |
| Lys3 | 8.08 | 124.01 | 4.32 | 56.00 | 1.71, 1.66 | 33.12 | Hγ 1.27, 1.33; Hδ 1.65; Hε 2.97; Cγ 23.96; Cδ 28.82; Cε 41.78 |
| Phe4 | 8.36 | 122.16 | 4.58 | 57.10 | 3.07, 3.17 | 39.24 | Hδ 7.41; Hε 7.35; Hζ 7.31; Cδ 131.16; Cε 131.48; Cζ 129.59 |
| C'$NH_2$ | 7.16,7.65 | 109.26 | | | | | |

---

[a]NMR data were collected on 10 mM peptide samples, at pH 6, and a temperature of 25 $^{o}$C. The samples contained no added buffers or salts. Chemical shifts were referenced using 3-(Trimethylsilyl)propane-1-sulfonate (DSS) and are given in units of ppm. Because the free and bicelle-bound peptides are in fast exchange, and since there is an excess of the peptides, the chemical shift assignments in this table are also valid for the bound peptides.
[b]Underlined $^{1}$H chemical shifts show significant differences of ~0.4 ppm from the random coil values reported in Table 2.3 of Wüthrich, K. NMR of Proteins and Nucleic Acids. (John Wiley & Sons, New York; 1986). These upfield shifts are characteristic of aromatic ring current effects, and likely reflect basic residues involved in cation-pi interactions with aromatic rings.

**Table S2. Statistics for the 20 lowest-energy NMR structures of peptides in their free states.**

| **NMR Structure** | *SPN10* | *SS20* | *SPN4* | *SS31* |
|---|---|---|---|---|
| **Restraints** | | | | |
| NOEs (total) | 39 | 52 | 46 | 29 |
| Intraresidue NOEs | 28 | 33 | 33 | 14 |
| Sequential NOEs | 11 | 19 | 10 | 12 |
| NOEs( \|i-j\| = 2) | 0 | 0 | 3 | 3 |
| NOEs (\|i-j\| = 3) | 0 | 0 | 0 | 0 |
| Dihedral ($\phi,\psi$)[a] | 4 | 4 | 4 | 4 |
| **Restraint Violations**[b,c] | | | | |
| NOE (Å) | 0.0825 ± 0.0049 | 0.0523 ± 0.0030 | 0.00026 ± 0.00028 | 0.0422 ± 0.0109 |
| Dihedral (°) | 0.06 ± 0.15 | 0.0 ± 0.0 | 0.62 ± 0.20 | 0.12 ± 0.48 |
| **RMSD Ideal Geometry** | | | | |
| Bonds (Å) | 0.0036 ± 0.0003 | 0.0036 ± 0.0004 | 0.0042 ± 0.0002 | 0.0015 ± 0.0004 |
| Angles (°) | 0.35 ± 0.02 | 0.40 ± 0.03 | 0.46 ± 0.02 | 0.31 ± 0.04 |
| Impropers (°) | 0.17 ± 0.02 | 0.28 ± 0.04 | 0.26 ± 0.04 | 0.16 ± 0.03 |
| **Ramachandran Statistics**[d] | | | | |
| Most favored (%) | 0 | 0 | 0 | 0 |
| Allowed (%) | 100 | 50 | 100 | 100 |
| Generously allowed (%) | 0 | 0 | 0 | 0 |
| Disallowed (%) | 0 | 50[e] | 0 | 0 |
| **Coordinate RMSD (Å)**[f] | | | | |
| Backbone (C$\alpha$+N+C'+O) | 1.32 ± 0.23 | 0.70 ± 0.15 | 0.56 ± 0.07 | 0.64 ± 0.24 |
| Heavy | 2.41 ± 0.59 | 1.62 ± 0.34 | 1.47 ± 0.34 | 1.16 ± 0.20 |

[a]Loose dihedral restraints of $\phi$ = -90 ± 70 and $\psi$ = +60 ± 120 deg were included for the central residues 2 and 3 of the tetrapeptides. For the SS20 peptide which has a D-arg at position 2, the $\phi$ restraint was set to +90 ± 70 deg.
[b]Values are given as the NMR ensemble mean ± s.d.
[c]Structures contained no distance violations greater 0.3 Å or dihedral violations greater than 5 degrees.
[d]Dihedral angles are only defined for the central residues 2 and 3 of the tetrapeptides.
[e]The SS20 peptide has a D-arg at position 2, and its dihedral angles occur in the left-handed region of Ramachandran space.
[f]RMSD values were calculated using the GROMACS *gmx rms* function.

**Table S3. Statistics for the 20 lowest-energy NMR structures of peptides in their bicelle-bound state.**

| **NMR Structure** | *SPN10* | *SS20* | *SPN4* | *SS31* |
|---|---|---|---|---|
| **Restraints** | | | | |
| NOEs (total) | 95 | 113 | 100 | 106 |
| Intraresidue NOEs | 37 | 56 | 46 | 35 |
| Sequential NOEs | 47 | 39 | 43 | 55 |
| NOEs( \|i-j\| = 2) | 7 | 3 | 11 | 16 |
| NOEs (\|i-j\| = 3) | 4 | 15 | 0 | 0 |
| Dihedral ($\phi,\psi$)[a] | 4 | 4 | 4 | 4 |
| **Restraint Violations**[b,c] | | | | |
| NOE (Å) | 0.0720 ± 0.0018 | 0.0896 ± 0.0010 | 0.0697 ± 0.0019 | 0.0825 ± 0.003 |
| Dihedral (°) | 0.76 ± 0.36 | 1.76 ± 0.18 | 0.63 ± 0.19 | 1.14 ± 0.23 |
| **RMSD Ideal Geometry** | | | | |
| Bonds (Å) | 0.0078 ± 0.0002 | 0.0084 ± 0.0001 | 0.0056 ± 0.0002 | 0.0106±0.0002 |
| Angles (°) | 0.97 ± 0.01 | 0.89 ± 0.02 | 0.56 ± 0.02 | 0.90 ± 0.03 |
| Impropers (°) | 0.65 ± 0.06 | 0.71 ± 0.03 | 0.34 ± 0.02 | 0.57 ± 0.04 |
| **Ramachandran Statistics**[d] | | | | |
| Most favored (%) | 0 | 0 | 50 | 50 |
| Allowed (%) | 100 | 50 | 0 | 50 |
| Generously allowed (%) | 0 | 50[e] | 50 | 0 |
| Disallowed (%) | 0 | 0 | 0 | 0 |
| **Coordinate RMSD (Å)**[f] | | | | |
| Backbone (C$\alpha$+N+C'+O) | 0.06 ± 0.02 | 0.07 ± 0.15 | 0.04 ± 0.02 | 0.11 ± 0.03 |
| Heavy | 0.47 ± 0.09 | 0.55 ± 0.10 | 0.71 ± 0.19 | 0.90 ± 0.24 |

[a]Loose dihedral restraints of $\phi$ = -90 ± 70 and $\psi$ = +60 ± 120 deg were included for the central residues 2 and 3 of the tetrapeptides. For the SS20 peptide which has a D-arg at position 2, the $\phi$ restraint was set to +90 ± 70 deg.
[b]Values are given as the NMR ensemble mean ± s.d.
[c]Structures contained no distance violations greater 0.3 Å or dihedral violations greater than 5 degrees.
[d]Dihedral angles are only defined for the central residues 2 and 3 of the tetrapeptides.
[e]The SS20 peptide has a D-arg at position 2, and its dihedral angles occur in the left-handed region of Ramachandran space.
[f]RMSD values were calculated using the GROMACS *gmx rms* function.

**Table S4. RMSD and $R_g$ values for NMR measurements and MD simulations.**

| Ensemble Source | Heavy Atom RMSD (Å) | | | |
|---|---|---|---|---|
| | SS-31 | SS-20 | SPN4 | SPN10 |
| NMR: Solution | 1.16±0.20 | 1.62±0.34 | 1.47±0.34 | 2.41±0.59 |
| NMR: Membrane | 0.90±0.24 | 0.55±0.10 | 0.71±0.19 | 0.47±0.09 |
| MD: Solution | 2.39±0.43 | 2.60±0.15 | 2.22±0.30 | 3.23±0.12 |
| MD Membrane | 1.89±0.27 | 1.89±0.43 | 2.04±0.34 | 1.83±0.42 |
| MD Membrane (restrained) | 1.93±0.28 | 2.07±0.34 | 2.03±0.29 | 1.44±0.37 |

| Ensemble Source | Heavy Atom Radius of Gyration (Å) | | | |
|---|---|---|---|---|
| | SS-31 | SS-20 | SPN4 | SPN10 |
| NMR: Solution | 4.57±0.12 | 4.70±0.13 | 4.72±0.16 | 4.79±0.20 |
| NMR: Membrane | 4.63±0.18 | 4.20±0.03 | 4.08±0.08 | 4.23±0.03 |
| MD: Solution | 5.38±0.13 | 5.42±0.08 | 5.44±0.05 | 5.22±0.13 |
| MD Membrane | 5.78±0.12 | 5.22±0.10 | 5.67±0.14 | 5.45±0.17 |
| MD Membrane (restrained) | 5.49±0.10 | 4.86±0.12 | 5.43±0.12 | 5.08±0.21 |

**Table S5. Improved structural agreement between MD and NMR when imposing NOE restraints.**

| Peptide Analog | % Time <3Å Heavy Atom RMSD | | | |
|---|---|---|---|---|
| | Unrestrained | | w/ NOE Restraints | |
| SS-31 | **Avg** | 5.70% | **Avg** | 52.11% |
| | **95% CI** | 3.43 - 7.97% | **95% CI** | 40.94 - 63.29% |
| | **Range** | 0.89 - 17.78% | **Range** | 5.24 - 95.11% |
| SS-20 | **Avg** | 2.90% | **Avg** | 20.27% |
| | **95% CI** | 1.30 - 4.50% | **95% CI** | 7.58 - 32.96% |
| | **Range** | 0.00 - 9.84% | **Range** | 0.01 - 94.79% |
| SPN4 | **Avg** | 1.95% | **Avg** | 57.98% |
| | **95% CI** | 0.89 - 3.01% | **95% CI** | 49.23 - 66.74% |
| | **Range** | 0.19 - 11.28% | **Range** | 7.93 - 92.13% |
| SPN10 | **Avg** | 0.01% | **Avg** | 32.10% |
| | **95% CI** | 0.00 - 0.01% | **95% CI** | 16.59 - 47.61% |
| | **Range** | 0.00 - 0.06% | **Range** | 0.00 - 97.66% |
